# Gut microbiota-dependent increase in phenylacetic acid induces endothelial cell senescence during aging

**DOI:** 10.1101/2023.11.17.567594

**Authors:** Seyed Soheil Saeedi Saravi, Benoit Pugin, Florentin Constancias, Aurélien Thomas, Sylvain Le Gludic, Meret Sarah Allemann, Gergely Karsai, Pratintip Lee, Cristina Menni, Ilias Attaye, Jürg H. Beer

**Affiliations:** Center for Translational and Experimental Cardiology, Department of Cardiology, University Hospital Zurich, University of Zurich, 8952 Schlieren, Switzerland; Laboratory of Food Biotechnology, Institute of Food, Nutrition and Health, Department of Health Sciences and Technology, ETH Zurich, 8092 Zurich, Switzerland; Faculty Unit of Toxicology, University Center of Legal Medicine, Lausanne University Hospital and University of Lausanne, Lausanne, Switzerland; Unit of Forensic Toxicology and Chemistry, University Center of Legal Medicine, Lausanne University Hospital and University of Lausanne, Geneva University Hospital and University of Geneva, Lausanne, Geneva, Switzerland; Center for Molecular Cardiology, University of Zurich, 8952 Schlieren, Switzerland; Department of Internal Medicine, Cantonal Hospital Baden, 5404 Baden, Switzerland; Institute of Clinical Chemistry, University Hospital Zurich, 8952 Schlieren, Switzerland; Department of Twin Research, King’s College London, St Thomas’ Hospital Campus, London SE1 7EH, UK; Amsterdam Cardiovascular Sciences, Diabetes & Metabolism, Amsterdam, Netherlands

**Keywords:** Aging, Gut microbiota, Endothelial senescence, Phenylacetic acid, Hydrogen peroxide, Epigenetic alteration, SASP

## Abstract

Endothelial cell (EC) senescence plays a crucial role in the development of cardiovascular diseases in aging population. Gut microbiota alterations are emerging as significant factors present in cellular senescence associated with aging. However, little is known about how aging-related changes in gut microbiota are causally implicated in EC senescence. Here we show that gut microbiota-dependent phenylacetic acid (PAA) and its derivative, phenylacetylglutamine (PAGln), are elevated in a human aging cohort (TwinsUK, n=7,303) and in aged mice. Metagenomic analyses revealed a marked increase in the abundance of PAA-producing microbial pathways (PPFOR and VOR), which were positively associated with the abundance of *Clostridium* sp. ASF356, higher circulating PAA concentrations, and endothelial dysfunction in old mice. We found that PAA potently induces EC senescence and attenuates angiogenesis. Mechanistically, PAA increases mitochondrial H_2_O_2_ generation, which aggravates IL6-mediated HDAC4 translocation and thereby upregulates VCAM1. In contrast, exogenous acetate, which was reduced in old mice, rescues the PAA-induced EC senescence and restores angiogenic capacity through markedly alleviating the SASP and epigenetic alteration. Our studies provide direct evidence of PAA-mediated crosstalk between aging gut microbiota and EC senescence and suggest a microbiota-based therapy for promoting healthy aging.

**Highlights:**

- Aging-related gut microbiota alterations contribute to a marked elevation of plasma PAA and PAGln in humans and mice
- *Clostridium* sp. ASF356 contributes to PPFOR-mediated PAA formation in aged mice
- Gut-derived PAA promotes endothelial senescence and impairs angiogenesis
- PAA induces mitochondrial H_2_O_2_ generation, by which drives epigenetic alterations and SASP in ECs
- Acetate rescues PAA-induced EC senescence and mitochondrial dysfunction
- Acetate improves angiogenesis by reducing HDAC4 phosphorylation and SASP

## Introduction

Aging is a significant risk factor for prevalent diseases including cardiovascular diseases (CVD), affecting millions of people worldwide^1^. The alarming increase in incident CVD in older adults emphasizes our inadequate understanding of the development of CVD in aging^2^. Dysfunctional endothelial cells (ECs) are increasingly recognized to contribute to CVD^3^. Advanced age causes cellular senescence in ECs, which is crucially involved in EC dysfunction in older adults^4^. Senescent ECs (SECs) undergo a variety of persistent alterations, particularly including epigenetic modifications, irreversible cell-cycle arrest, and DNA instability, and produce senescence-associated secretory phenotype (SASP) that confer deleterious effects on angiogenic processes and vascular function^2,5^. Therefore, accumulation of SECs is presumed to be a primary contributor to aging-related CVD.

Multiple initiating triggers, namely oxidative stress, have been identified to drive stress-induced senescence in ECs^6,7^. Converging evidence supports the notion that gut microbiota, a yet underappreciated endocrine organ, can regulate oxidative-regulated host genes involved in endothelial cell biology^8^. Although such studies have helped pinpointing the association between gut microbial community and CVD^9,10^, there is limited evidence on gut microbiota-EC senescence crosstalk and its molecular basis in aging organisms.

With advanced age, alterations of specific gut-derived metabolites such as aromatic amino acid derivatives might contribute to CVD through extended oxidative stress and inflammatory response^11^. One such example is the accumulation of uraemic toxins that lead to accelerated aging of the cardiovascular system^11^. The newly described phenylacetic acid (PAA) and its derivative phenylacetylglutamine (PAGln) produced from phenylalanine by commensal microbial *porA* or *ppfor* genes^12^ have been positively associated with cardiovascular and all-cause mortality in patients with chronic kidney disease (CKD)^13^ and heart failure^14^. In preclinical models, PAA stimulated the production of reactive oxygen species (ROS) and further inflammation in aortic endothelial cells^15^, explaining its role in pathological changes of the vasculature in CKD. Similarly, through pathways involving ROS production and apoptosis, PAA-derivative PAGln fostered cardiomyocyte damage and decreased contractility^14,16^. In this study, we first observed higher plasma concentrations of PAA and PAGln in both humans enrolled in the TwinsUK aging cohort and old mice. We, therefore, hypothesized that elevated plasma levels of these two metabolites during aging may actively promote EC senescence, leading to reduced angiogenesis and vascular dysfunction. We aimed to uncover mechanisms underpinning the crosstalk between aging gut-derived PAA and EC senescence and dysfunction. Moreover, restoration of angiogenic capacity in these cells is a therapeutic objective.

Beyond the metabolites, gut microbiota produces a myriad of other metabolites such as short-chain fatty acids (SCFAs) mainly including acetate, propionate and butyrate which circumvent atherosclerosis and other CVD by tackling multiple risk factors^11^. It is well defined that >70% of colonic acetate, but only small amounts of propionate and butyrate, reach the circulation after hepatic metabolism^17^. We have previously reported that aging markedly decreases fecal acetate levels up to 80% by reducing some bacteria, for instance *Prevotella* and *Rikenellaceae_RC9_gut_group*, which contribute to production of colonic acetate^18^. The depletion of acetate producers in the gut is observed ubiquitously in CVD and tracks with disease severity^19^. Converting acetate to acetyl-CoA in mitochondria, quiescent ECs which are sensitive to oxidative stress utilize acetyl-CoA to sustain the tricarboxylic acid (TCA) cycle and redox homeostasis and to protect themselves against dysfunction^20^. Acetate also controls epigenetic alterations by regulating histone acetylation/deacetylation critical to endothelial cell phenotype and function^20^. Accordingly, we hypothesized that exogenous acetate might suppress PAA-induced oxidative stress and associated epigenetic alterations driving EC senescence with therapeutic relevance to counteract EC dysfunction during aging. Interestingly, our findings revealed a new role for acetate in EC senescence escape and provide new insights into how acetate can protect against age-related arterial dysfunction and associated CVD.

## Results

### Gut Microbiota is associated with age-dependent elevation of plasma PAA and PAGln in humans and mice

Our data identified markedly higher concentrations of PAA and PAGln in plasma obtained from old mice (>24-months) compared to young mice (>3-months), as detected by targeted metabolomics (Fig. 1a-c). Of note, studies were performed in old and young mice with relatively preserved kidney function, indicated by similar serum BUN and creatinine levels (*p*>0.05; Supplementary Fig. 1a,b), to exclude the impact of age-related kidney failure on the elevated plasma levelsof PAA and PAGln in old mice. The mouse groups were also matched by sex and body weight (Supplementary Fig. 1c). Similar findings were observed in blood samples derived from the high-resolution gut microbiome–plasma metabolome study from the TwinsUK adult twin registry^21^. Non-targeted metabolomics analysis of plasma from the human aging cohort reflected the age-associated concentration changes of both PAA (Fig. 1d) and PAGln (Fig. 1e) as seen in mice. Accordingly, our linear regression analysis between metabolite concentrations and age revealed that PAA and PAGln age-dependently increase in 7,303 individuals between 18 and 95 years old (*r* = 0.06, *R^2^* = 0.003, *p*<0.001 for PAA; *r* = 0.25, *R^2^* = 0.063, *p* <0.001 for PAGln).

**Fig. 1.**
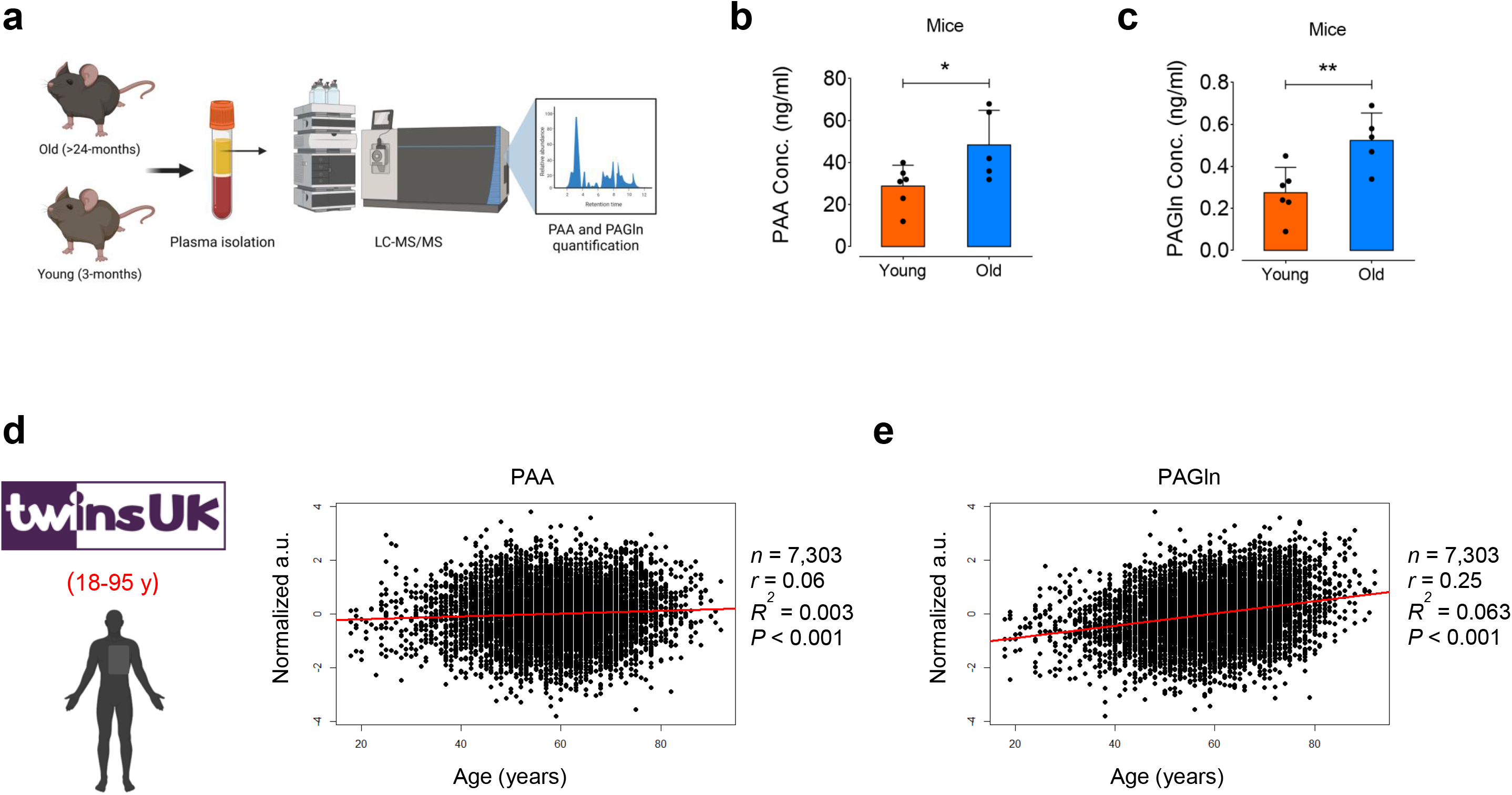
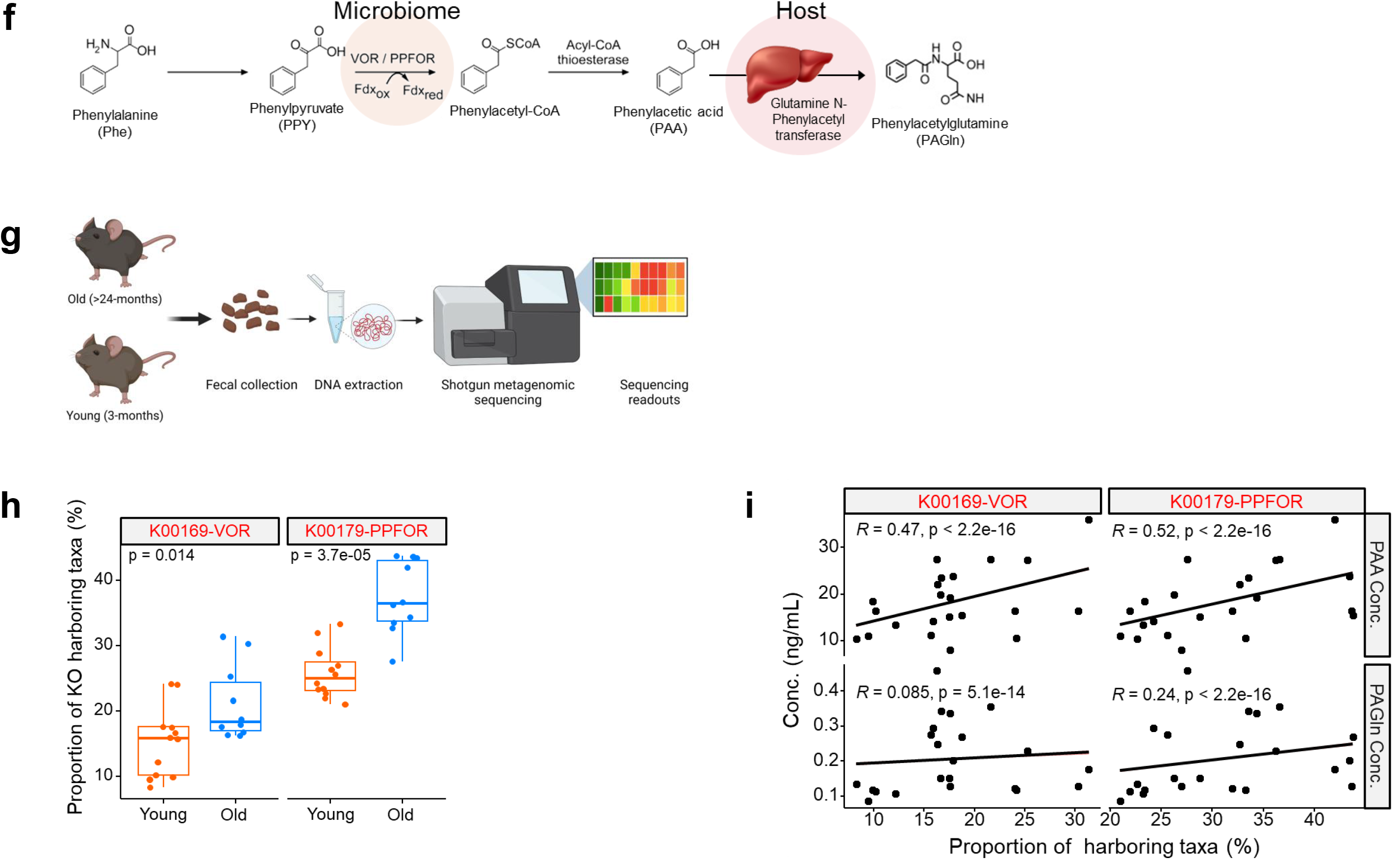
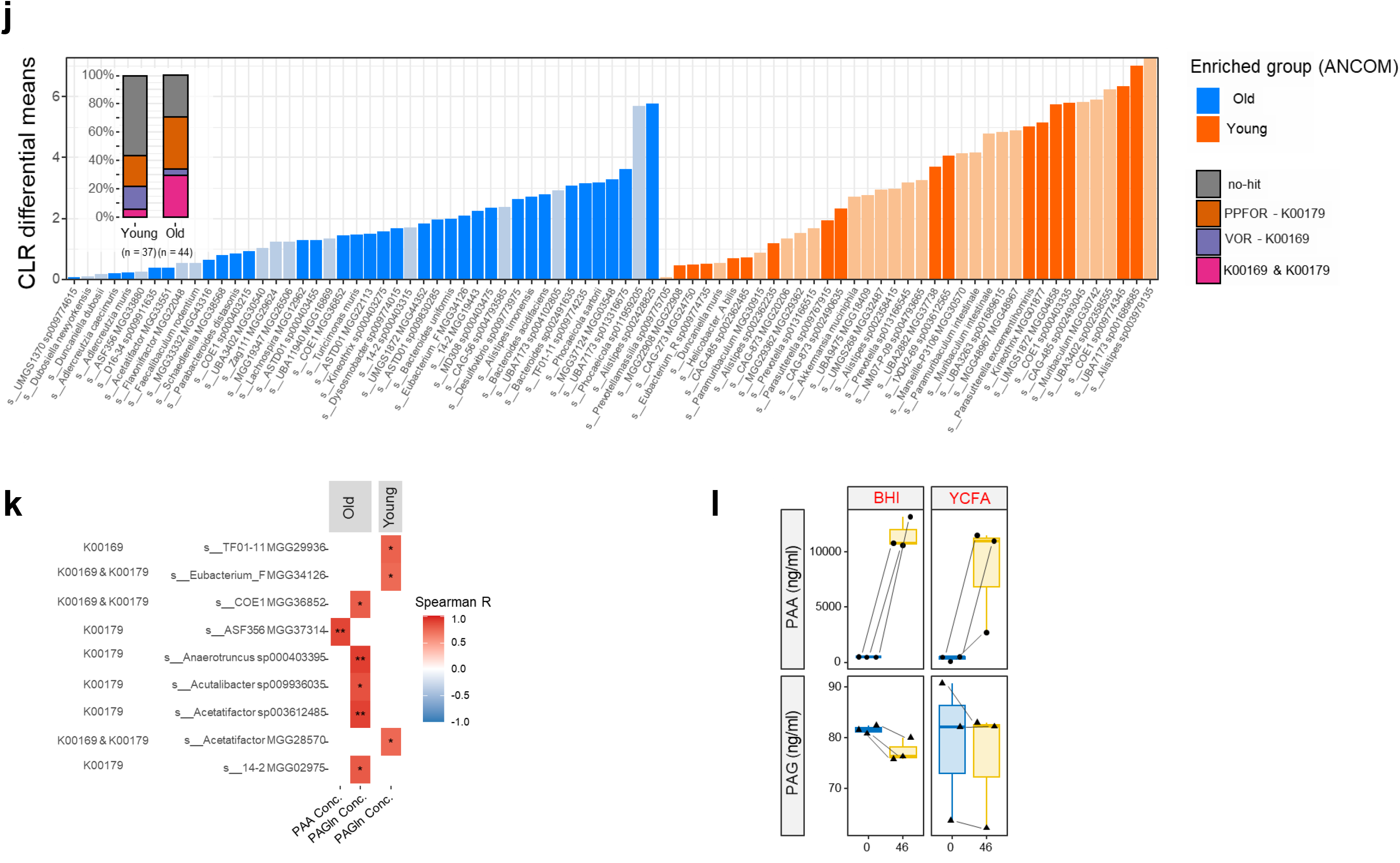
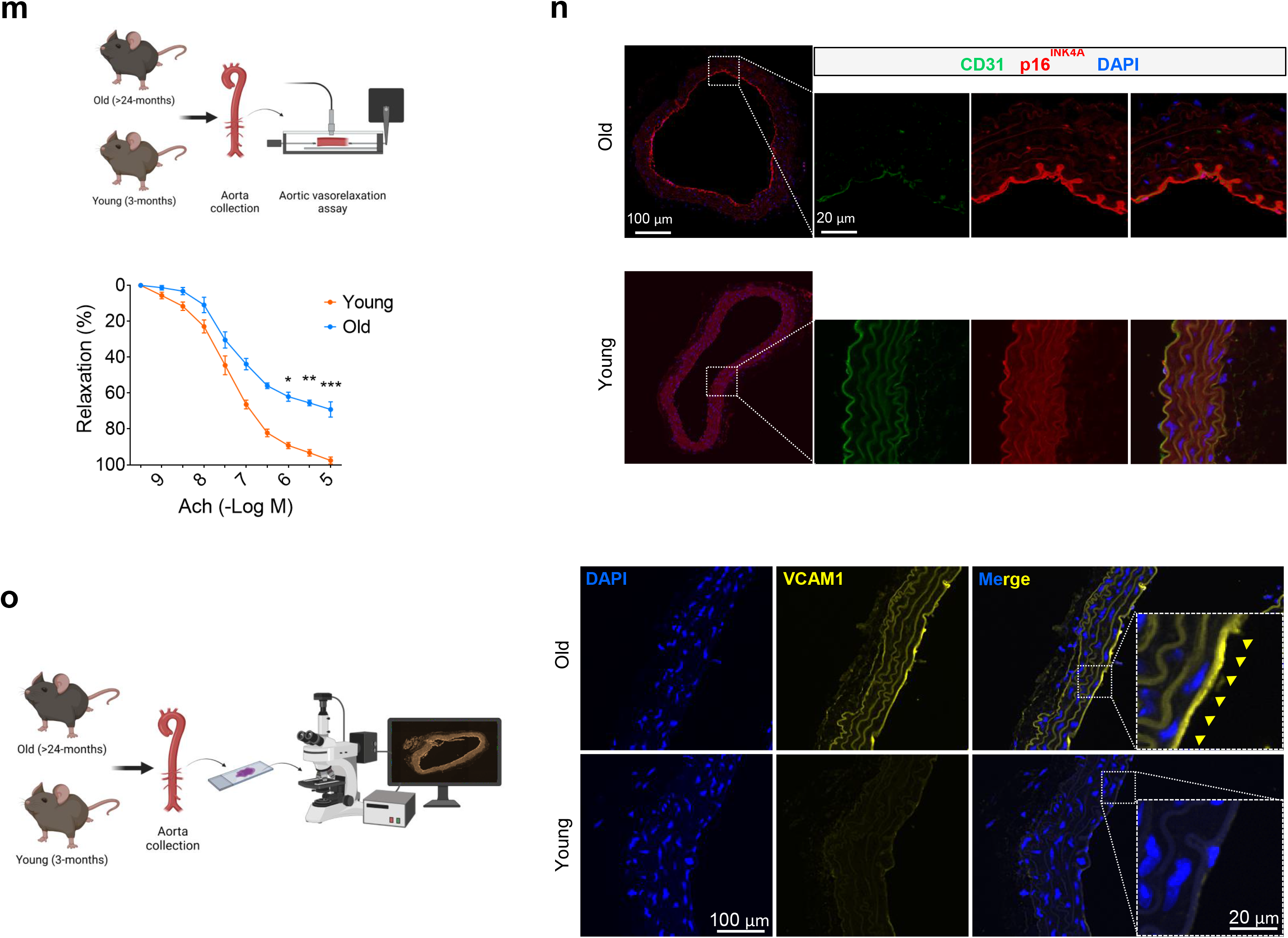
Reflection of age in plasma levels of gut microbiota-derived metabolites PAA and PAGln. **a,** Schematic diagram of the experimental setting: >24 (old) and 3 (young)-month-old C57BL/6J male mice were subjected to LC-MS/MS quantification of plasma PAA and PAGln. **b,c,** PAA (**b**) and PAGln (**c**) levels were significantly higher in plasma of old mice compared to young mice (n=6). **d,e,** Dot plots of PAA (**d**) and PAGln (**e**) quantified by non-targeted metabolomics on plasma demonstrates a positive association with age in the TwinsUK cohort (n=7,303). r: Correlation coefficient. **f**, Schema for the metaorganismal production of PAA from dietary Phenylalanine (Phe) by the VOR/PPFOR-mediated system. **g,** Schematic diagram of the experimental process of shotgun metagenomics of feces from old and young mice. **h,** Distribution of VOR and PPFOR homologs in mouse microbiome genomes. The box plots show the higher proportion (%) of microbial genomes harboring VOR and PPFOR homologs in old mice compared to young mice (n=5-6). Analyses were performed using the KEGG orthology (KO) functions; KOO169 (for VOR) and KOO179 (for PPFOR). **i,** Gene–metabolite interactions explain a positive correlation of the proportion (%) of either VOR- or PPFOR-harboring taxa and plasma PAA levels in old mice. Nevertheless, the lowermost chart shows non-significant association of VOR or PPFOR homologs and plasma PAGln levels in these mice (n=5-6). **j,** Bar plot depicting the age-associated relative abundance of fecal microbiota profiles at the strain level (Dark colors represent the bacterial taxa harboring VOR or PPFOR homologs). The leftmost bar plot demonstrates the proportion of VOR, PPFOR, or VOR+PPFOR homologs detected in taxa enriched in old (n=44) *versus* young (n=37) mice. **k,** Heatmap of Spearman correlations denotes a positive association between plasma levels of PAA or PAGln and representative gut bacteria among the top differentially abundant microbial taxa in old *versus* young mice. **l,** PAA is produced anaerobically from Phe by *Clostridium* sp. ASF356 cultured in both BHI (left) and YCFA (right) media (top). No significant changes were seen in PAGln levels in the supernatants obtained from the bacterial culture incubated with Phe (Lower panel). **m,** Schematic diagram of the experimental setting: Aortas were harvested from old and young mice for *ex vivo* force tension myography. Vasorelaxation responses (%) *ex vivo* to acetylcholine (Ach) in aortic rings from old mice were significantly lower than young mice (n=10). **n,** P16^INK4a^ immunofluorescence staining reveals a hallmark of cellular senescence in CD31-positive ECs in the ascending aortas of old mice (n=5-6). **o,** VCAM1 immunofluorescence demonstrates higher expression of VCAM1 in the ascending aortic endothelial cells from old mice compared to those from young mice (n=5-6). Scale bars, 20 and 100 μm. Error bars represent SD (**b,c**) or SEM (**h**,**i,l,m**) or 95% confidence intervals (**d,e**). Continuous data are presented as mean ± SD or SEM. Statistical analysis was performed with a two-tailed unpaired Student’s *t*-test (**b,c,h,l,m**), a two-tailed test for Pearson correlation analysis (**d,e**), partial Spearman’s correlation test (**i,k**), and ANCOM method (**j**). (\**P*<0.05, \*\**P*<0.01, \*\*\**P*<0.001). Source data are provided as a Source Data file.

Previous studies have shown that in both humans and mice, the gut microbiota can convert dietary phenylalanine into PAA via a metaorganismal pathway, at which point host PAA conjugation with glutamine leads to PAGln generation^12^. Accordingly, antibiotic suppression of gut microbiota had revealed a marked suppression of circulating PAGln levels^12,22^. These data emphasize that production of PAA and its byproduct PAGln is a gut microbiota-dependent process. As previously shown, two distinct gut microbial pathways, one catalyzed by phenylpyruvate:ferredoxin oxidoreductase (PPFOR) and the other one by α-ketoisovalerate:ferredoxin oxidoreductase (VOR), are involved in PAA production via oxidative decarboxylation of phenylalanine^12^ (Fig. 1f). In order to investigate the relationship between PPFOR and VOR gene abundances and aging, we performed genome shotgun metagenomic analyses of fecal samples from old and young mice (Fig. 1g). Using the Kyoto Encyclopedia of Genes and Genomes (KEGG) database, we identified a significantly higher abundance of both PPFOR and VOR pathways in gut microbiome of old *versus* young mice (*p*=3.7e^-5^ for PPFOR and *p*=0.014 for VOR; Fig. 1h). In addition, we observed a significant correlation between plasma PAA levels and the percentage of either PPFOR- or VOR-positive genomes (*R^2^*=0.52 for PPFOR and *R^2^*=0.47 for VOR, p<0.0001; Fig. 1i), highlighting a major role for gut microbiota in PAA production *in vivo* in mice. However, a weaker correlation between plasma PAGln concentrations and percentage of VOR-harboring bacteria were seen in the microbiome of old mice (*R^2^*=0.24 for PPFOR and *R^2^*=0.085 for VOR, p<0.0001; Fig. 1i). These suggest that age-related higher abundance of both PPFOR and VOR pathways is positively associated with plasma levels of PAA and PAGln metabolites.

Our metagenomic analyses revealed significant alterations in both alpha-diversity indices of richness (Hil-d0, 1, and 2) and evenness (evenness_pielou) (Supplementary Fig. 2a) as well as beta-diversity (principal coordinate analysis with Aitchison distance; Supplementary Fig. 2b) in the gut microbiome in old mice as opposed to young mice. In the line with the above findings, the centered log-ratio (CLR) by analysis of composition of microbiomes (ANCOM) demonstrates differentially abundant bacterial taxa harboring VOR, PPFOR, or both homologs in the microbiome of old compared to young mice. As shown in Fig. 1j, the number of dark bars representing VOR- or PPFOR-harboring taxa in the old microbiome is markedly higher than in young mice. Accordingly, ∼1.7-fold higher proportion of VOR, PPFOR, or VOR+PPFOR homologs were detected in taxa enriched in old mice compared to those in young mice (72% vs. 42%; Fig. 1j).

We next aimed to identify the specific PPFOR- or VOR-harboring bacteria which are associated with increased circulating PAA or PAGln levels in the old host. Our Spearman’s rank correlation analysis shows that five taxa (including four PPFOR-harboring *Anaerotruncus sp000403395*, *Acutalibacter MGG29203*, *Acetatifactor sp003612485*, *Lachnospiraceae 14-2 MGG02975*, and one VOR+PPFOR-harboring *Lachnospiraceae COE1 MGG36852*, all from class *Clostridia*) were positively correlated with plasma PAGln in old mice and only one taxa (*Clostridium ASF356 MGG37314*) was positively correlated with PAA levels in plasma obtained from old mice (Fig. 1k).

To further confirm the functional role of *Clostridium ASF356 MGG37314* in PAA formation from phenylalanine, we cultivated axenic culture of *Clostridium* sp. ASF356 and tested for PAA production by mass spectrometry (LC-MS/MS). We observed that *Clostridium* sp. ASF356 harboring PPFOR gene homolog produced PAA when grown in phenylalanine-containing BHI or YFCA media under anaerobic condition (Fig. 1l), while no changes in PAGln concentrations were detected (Fig. 1I). These results suggest that *Clostridium* sp. ASF356 commensal in the aged microbiota contributes to increased circulating PAA levels in old mice.

On the other hand, we observed a markedly disrupted endothelial function in old mice, as reflected by decreased endothelium-dependent relaxation to acetylcholine, compared to what seen in young mice. As shown in Fig. 1m, there was a significantly reduced maximal relaxation at 10^−5^LM (*p*L<L0.001; Young: 97.57L±L1.91; Old: 69.13L±L4.25), and rightward shift of dose-response curve to acetylcholine (*p*L<L0.05; EC_50_, Young: 34.23L±L9.01 nM; Old: 51.33L±L12.32 nM) was seen in isolated aortas from young *versus* old mice. The aging-associated endothelial dysfunction was accompanied by a remarkable abundance of p16INK4A, as a hallmark of cellular senescence, in CD31^+^ aortic ECs from old mice as opposed to young mice (Fig. 1n). Additionally, aortic endothelial cells from old mice exhibit a higher expression of VCAM1 as compared to those in young aortas (Fig. 1o).

The positive association between plasma levels of PAA and PAGln metabolites with age in mice (and similarly in humans) alongside concurrent vascular dysfunction, underscores the potential key role of phenylalanine-derived metabolites in the alteration of EC biology and vascular decline in aging.

#### PAA Directly Promotes Premature Senescence in ECs

To analyze the role of PAA in EC senescence, we cultured proliferating human aortic endothelial cells (passage 4-5) and stimulated them with PAA to test hallmarks of cellular senescence. We prepared replicative senescent HAECs (passage 15-17), as control. The senescence phenotype (such as enlarged, flattened, and multinucleated appearance of cells) and associated markers were validated through reduced proliferation and increased SA-β-galactosidase activity, telomere shortening, and expression of CDK inhibitors and SASP factors in proliferating ECs (PEC) *versus* PAA-exposed PECs *versus* replicative senescent ECs (SEC) (Fig. 2a, Supplementary Fig. 3). When treated with PAA, proliferating ECs exhibited multiple premature senescence-like features such as increased numbers of SA-β-gal^+^ cells (Fig. 2b) and γ-H2A.X immunofluorescence comparable to what seen in SECs (p.15-17) (Fig. 2c). We next sought to determine how PAA contributes to cell-cycle arrest in PECs. As a result, PAA-dependent senescence is accompanied by a concomitant increase in expression of CDK inhibitors p16^INK4a^, p19^INK4d^, and p21^WAF1/Cip1^ at mRNA level, as seen in SEC at passage 16 (Fig. 2d).

**Fig. 2.**
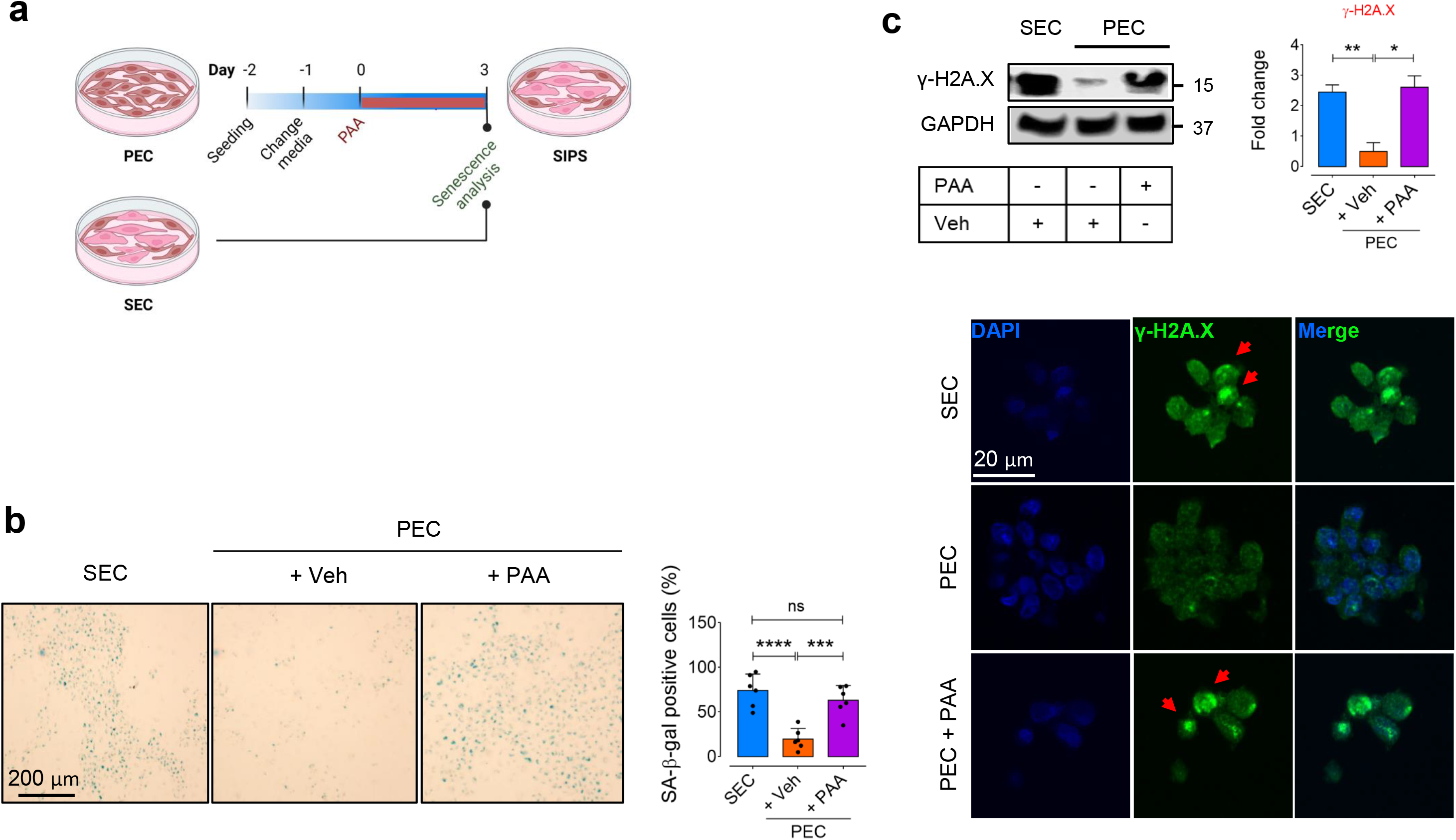
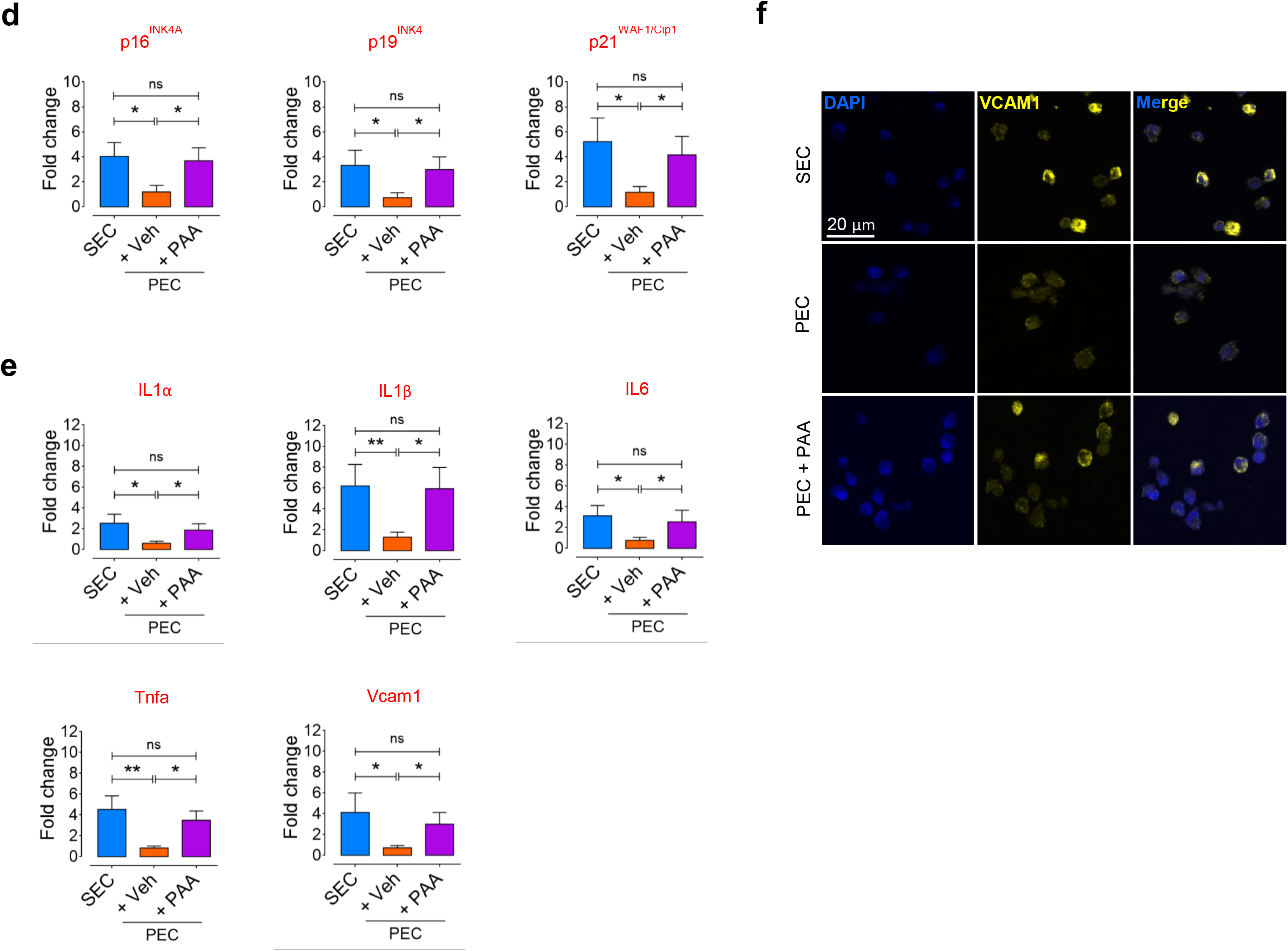
PAA induces endothelial cell senescence. **a,** Schematic diagram of the experimental setting: μμProliferating ECs (PEC) at p4-5 were treated with PAA (10 μM) for 72 h and then subjected to senescence hallmarks profiling. **b,** SA-β-gal staining shows an increase in cellular senescence in PECs treated with PAA at the magnitude seen in replicative senescent ECs (SEC) at p15-17. Right, quantitative plots are shown for SA-β-gal-positive cells (%) (n=6). **c,** qPCR shows that PAA treatment resulted in increased expression of CDK inhibitors p16^INK4a^, p19^INK4d^, and p21^WAF1/Cip1^ in PECs. **d,** Immunoblots (top) and γ-H2A.X immunostaining (bottom) represent DDR in PAA-exposed PECs compared to vehicle-treated PECs (n=6). **e,** qPCR demonstrates that SASP genes IL1α, IL-1β, IL-6, Tnfa, and VCAM1 are significantly upregulated in PAA-treated PECs (n=6). **f,** VCAM1 immunostaining shows its marked overexpression in PAA-induced senescent ECs (n=6). Data from *in vitro* cellular experiments represent triplicated biologically independent experiments. Scale bar, 20 μm. Error bars represent SD (**b-e**). *P* values were calculated using one-way ANOVA followed by Tukey’s post *hoc* test (**b-e**). (\**P*<0.05, \*\**P*<0.01, \*\*\**P*<0.001, \*\*\*\**P*<0.0001, ns, not significant). Source data are provided as a Source Data file.

In addition, using a qPCR-based assay, we also detected various SASP markers expression in PAA-treated PEC versus PEC, suggesting a positive correlation between PAA and the upregulation of SASP genes, including *Il1*α, *Il1*β, *Il6*, *Tnf*α, and *Vcam1* (Fig. 2e). Our VCAM1 immunostaining also demonstrates an increase in VCAM1 expression in PEC exposed to PAA (Fig. 2f). These are consistent with relevant findings observed in Fig. 1o demonstrating markedly higher VCAM1 immunofluorescence in old *versus* young aortas. Together, these data support a potential role of PAA in the regulation of EC senescence.

#### PAA Stimulates Mitochondrial Dysfunction by Stimulating ROS Generation in ECs

We observed that PAA triggers EC premature senescence to the magnitude seen in exogenous H_2_O_2_-exposed PEC (Fig. 3a). Excessive ROS such as H_2_O_2_ are well characterized to trigger EC senescence and dysfunction^23^; we, therefore, tested whether PAA increases ROS generation. Accordingly, we demonstrated a significantly higher CellRox green fluorescence, which represents increased global ROS generation, in PAA-exposed PEC *versus* vehicle-treated PEC (Fig. 3b). Using the ultrasensitive H_2_O_2_ biosensor, HyPer 7.2, we detected a marked increase in H_2_O_2_ production in the mitochondria of PEC (exposed to PAA) (Fig. 3c). As shown in Fig. 3d, maximal HyPer7.2 ratio in PECs in the presence of PAA exhibited no significant difference to those in the presence of exogenous H_2_O_2_ (50 μM; *p*L<L0.01). To further explore the major source of mitochondrial H_2_O_2_ production, we found that NADPH oxidase 2 (NOX2) expression increases in the presence of PAA in PECs compared with vehicle-incubated PECs (Fig. 3e). Since glutathione peroxidase converts oxidized glutathione (GSSG) to its reduced form (GSH), a key cellular ROS-scavenger^24^, we measured GPx1 transcriptional levels in baseline conditions and upon exposure to exogenous PAA (as "stress condition"). Levels of *Gpx1* transcripts showed a significant reduction in PECs in stress condition induced by PAA, supporting a role of PAA in counteracting simultaneous antioxidant defense and inducing oxidative stress (Supplementary Fig. 4).

**Fig. 3.**
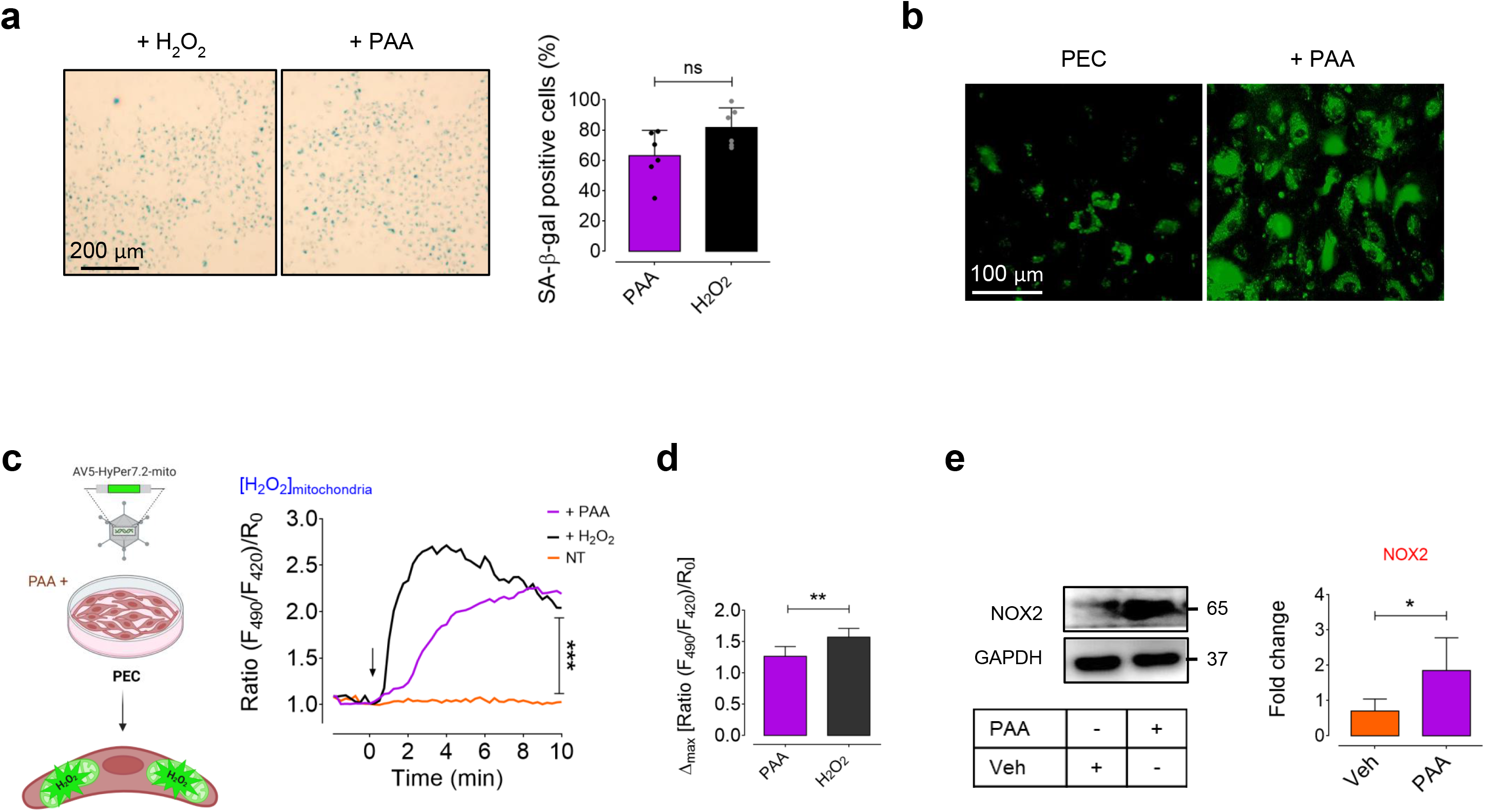
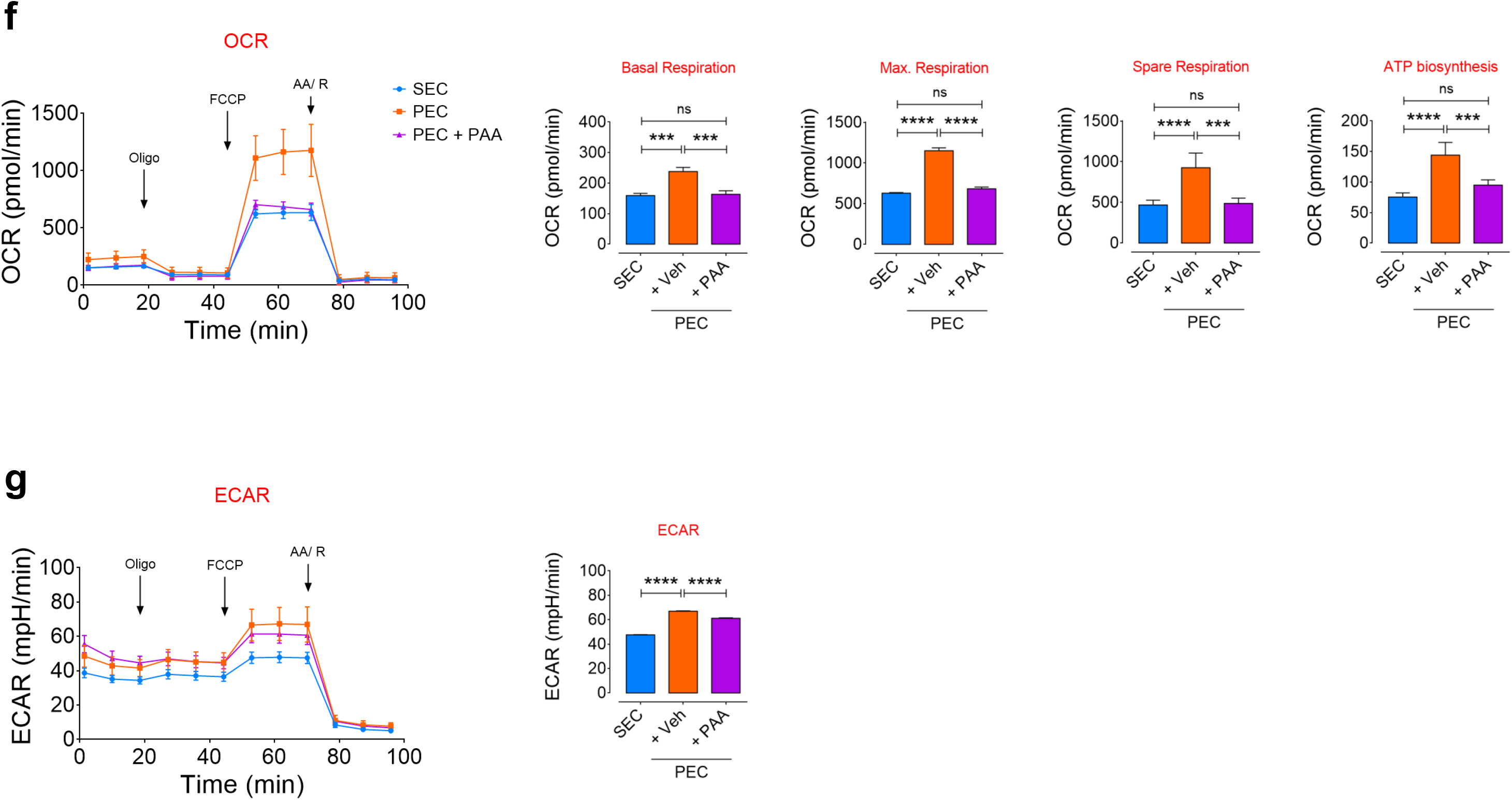
PAA regulates premature EC senescence by H_2_O_2_-mediated mitochondrial impairment. **a,** SA-β-gal staining shows that PAA induces cellular senescence in PECs comparable to the level seen in PECs incubated with exogenous H_2_O_2_ (50 μM). Quantitative plots for SA- β-gal-positive cells (%) are shown (n=6). **b,** CellRox green staining reveals that PECs markedly increase generation of intracellular ROS, including H_2_O_2_, (as green fluorescence) in the presence of PAA. **c,** Schematic diagram of the experimental setting: PECs were transduced with adenovirus 5 (AV5)-HyPer7.2 targeted to the cell mitochondria for further ratiometric fluorescence imaging to detect mitochondrial H_2_O_2_ generation in the presence of PAA. Right, the curves demonstrate significantly higher H_2_O_2_ responses in the mitochondria of HyPer7.2-transfected PECs following treatment with either PAA (n=25) or exogenous H_2_O_2_ (n=27) compared to vehicle (n=24). **d,** Bar chart shows maximal mitochondrial H_2_O_2_ responses to PAA or exogenous H_2_O_2_ (n=6). **e,** Immunoblot analysis demonstrates that PAA treatment results in a significant increase in expression of NOX2 in PECs (n=6). **f,** Mitochondrial respiratory rates in PECs treated with either PAA or vehicle and in replicative SECs was measured using Seahorse flux analyzer by Cell Mito Stress kit (n=6). Bar charts reveal significant reduction in oxygen consumption rate (OCR) represented by decreased maximal, basal, and spare reserve, and ATP biosynthesis in PAA-exposed PECs compared to PECs incubated with vehicle. **g,** Glycolysis was measured by extracellular acidification rates (ECAR) in vehicle- or PAA-treated PECs and replicative SECs under basal conditions (n=6). Right, Bar chart depicts a significantly higher ECAR in vehicle-treated PECs compared to PAA-exposed PECs or replicative SECs. Data from *in vitro* cellular experiments represent triplicated biologically independent experiments. Scale bars, 100 and 200 μm. Error bars represent SD (**a,d-g**) or SEM (**c,f,g**). *P* values were calculated using a two-tailed unpaired Student’s *t*-test (**a,e**), two-tailed Mann-Whitney *U*-test (**d**), and one-way ANOVA followed by Tukey’s post *hoc* test (**c,f,g**). (\**P*<0.05, \*\**P*<0.01, \*\*\**P*<0.001, \*\*\*\**P*<0.0001, ns, not significant). Source data are provided as a Source Data file.

As altered EC metabolism is a key driver of EC senescence^25^, we tested mitochondrial respiration, impairment of which generates oxidative stress, leading to EC dysfunction. In line with this, our standard Seahorse mitochondrial stress test in a high glucose medium displayed a decrease in basal, and more strikingly maximal, respiration rates (up to 2-fold) in PAA-induced senescent ECs (to the magnitude seen in SECs) compared with vehicle-incubated PECs (Fig 3f). Moreover, PECs exhibited a reduction in spare respiratory capacity, with FCCP-stimulated oxidative phosphorylation (OXPHOS) uncoupling, accompanied by decreased ATP biosynthesis in response to PAA (Fig. 3f). In agreement with the observed reduction in mitochondrial respiration, PAA-exposed PECs exhibited a markedly decreased basal extracellular acidification rate (ECAR), suggesting that the cells encounter a decline in glycolysis, which is a major energy supplier that generates ∼85% of total ATP in endothelial cells^26^ (Fig. 3g).

#### PAA-induced EC Senescence is mediated by Epigenetic Alterations

EC senescence is distinguished by deep epigenetic reprogramming, including histone modifications which elicit extensive changes of gene expression patterns that thus lead to vascular aging^4,27^. HDAC4 is an epigenetic regulator that critically controls cellular senescence^28,29^. Indeed, post-translational modification of HDAC4 contributes to cell proliferation, which is partly regulated by NOX4-derived H_2_O_2_ in endothelial cells^30^. In the line with this, our findings demonstrate that PAA induces mitochondrial H_2_O_2_ generation (Fig. 3b-d) which might regulate HDAC4 and epigenetic alterations, rendering EC senescence. We, therefore, thought to analyze the effects of PAA on epigenetic alterations in PECs.

Consistent with what seen in Fig. 2e, PAA significantly increases the expression of IL6 at protein level in ECs, while showed no differences compared to that in SECs (Fig. 4a,b). As seen in Supplementary Fig. 5a-f, DAAO-generated H_2_O_2_ (in presence of D-alanine) triggers the upregulation of IL6 in ECs. Because CaMKII is a both H_2_O_2_- and IL6-responsive protein kinase which regulates post-translational modification of HDAC4^30^, we assessed how PAA regulates CaMKII-HDAC4 pathway. Our findings exhibited a markedly increased phosphorylation of CaMKII at Thr286 and its downstream HDAC4 at Ser632 in PECs exposed to PAA (Fig. 4a,c,d). Using siRNA directed against CaMKII, protein expression was suppressed by ∼80% compared with negative siRNA-transfected cells for up to 3 days post-transfection (Supplementary Fig. 6a,b), leading to a demonstrable reduction in HDAC4^S632^ phosphorylation in PECs treated with PAA (Supplementary Fig. 6c). Increasing HDAC4 phosphorylation, PAA facilitates its nuclear export towards cell cytosol, hence we observed a markedly reduced expression of HDAC in the cell nucleus rather than cell cytosol in PAA-exposed PECs (Fig. 4i). Our immunostaining also confirms that PAA elicits cytosolic localization of HDAC4 (Fig. 4j), as seen in response to intracellular H_2_O_2_ generated by DAAO (Supplementary Fig. 7a,b). Moreover, an increased HDAC4 phosphorylation has been accompanied by a significant reduction in acetylation of histone 3 (H3) in response to PAA in PECs (Fig. 6a,e). Corroborating our findings, previous studies reveal that HDAC4 is post-transcriptionally downregulated in the nucleus of different cell types upon aging^31^. It exacerbates SASP and inflammatory signals and thereby induces cellular senescence^29^.

**Fig. 4.**
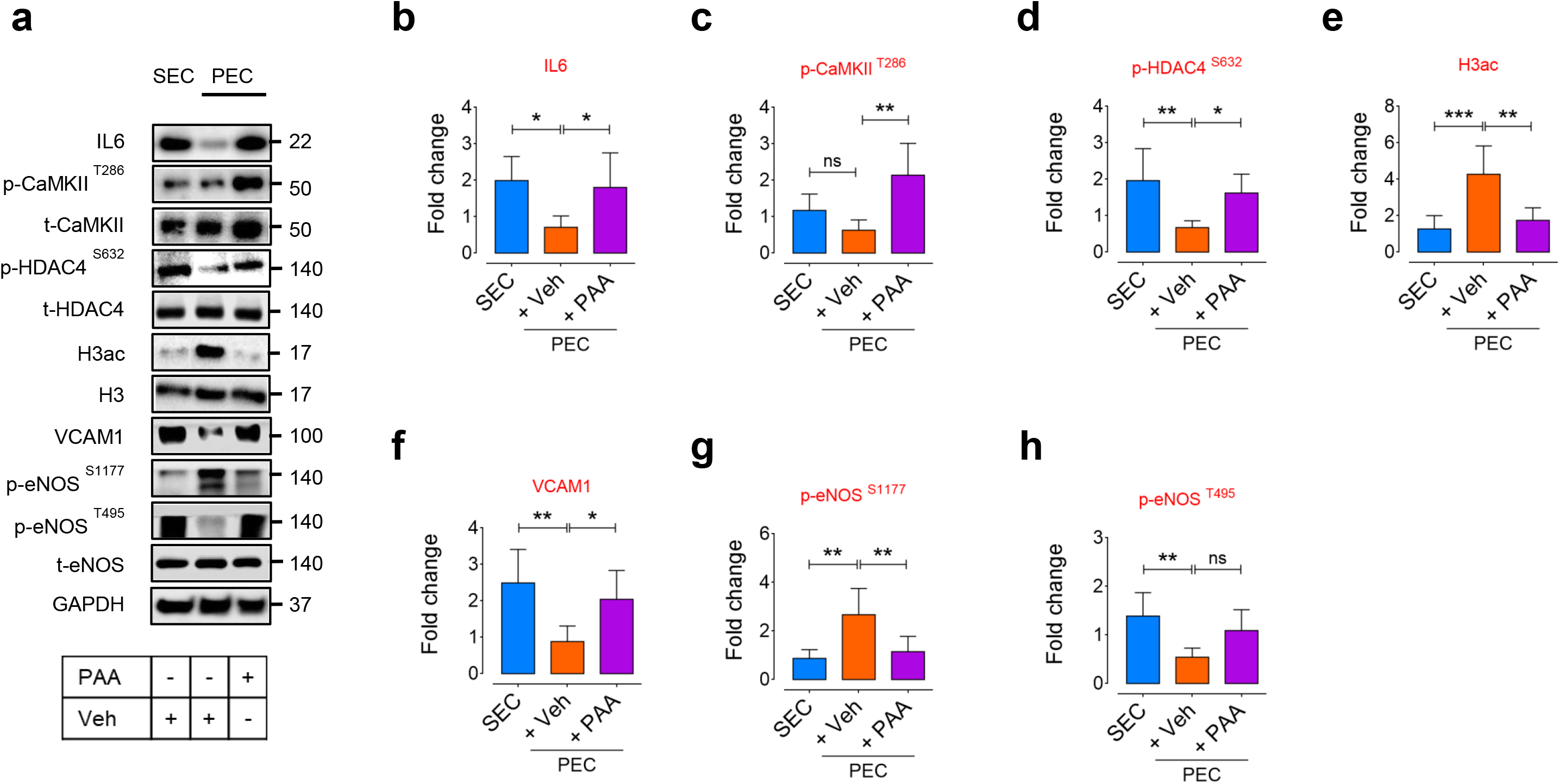
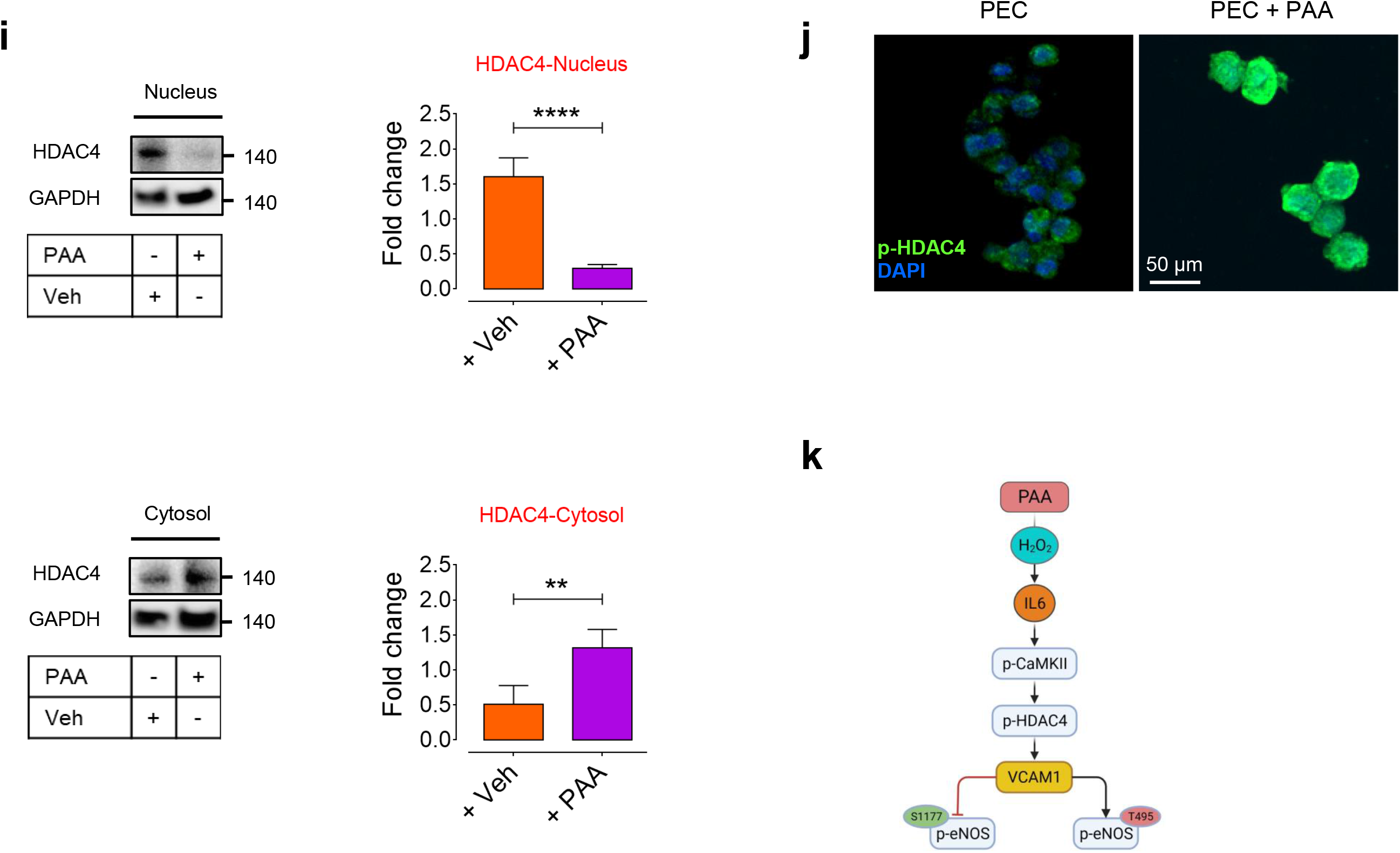
PAA induces EC senescence through the SASP-epigenetic regulation. **a-h,** Immunoblot analysis reveals the induction of IL6 (**b**) and CaMKII-mediated phosphorylation of HDAC4 at Ser632 (**c,d**) accompanied by decreased H3ac (**e**) in response to PAA in PECs. VCAM1 expression (**f**) was also induced by PAA, which was accompanied by post-transcriptional modifications of eNOS, represented as reduced phosphorylation at Ser1177 and increased phosphorylation at Thr495 (**g,h**) in PECs (n=6). **i,** Immunoblots (left) and quantitative plots (right) reveal translocation of HDAC4, represented as a decreased expression in the nucleus (top) and an increased expression in the cytosol (bottom) in PECs in response to PAA (n=6). **j,** p-HDAC4 immunostaining shows that PAA markedly increases HDAC4 phosphorylation that facilitates its translocation towards the cell cytosol in PECs (n=6). **k,** Summary scheme outlining the proposed signaling pathway by which PAA induces H_2_O_2_-mediated IL6 overexpression in proliferating endothelial cells. PAA interacts with CaMKII, which stimulates HDAC4 phosphorylation and subsequent nuclear export. It de-represses VCAM1 and reduces phosphorylation of its downstream regulator of endothelial function, eNOS, at Ser1177. Scale bar, 50 μm. Data from *in vitro* cellular experiments represent triplicated biologically independent experiments. Error bars represent SD (**b-i**). *P* values were calculated using one-way ANOVA followed by Tukey’s post *hoc* test (**b-h**) and a two-tailed unpaired Student’s *t*-test (**i**). (\**P*<0.05, \*\**P*<0.01, \*\*\**P*<0.001, ns, not significant). Source data are provided as a Source Data file.

One major component of HDAC4-directed regulation of cellular processes is the upregulation of VCAM1 in ECs^32^. We sought to determine if expression of VCAM1 was enhanced as a possible consequence of HDAC4 nuclear export by PAA. Consistent with an increase in *Vcam1* transcript by PAA treatment (Fig. 2e,f), we demonstrated increased VCAM1 protein expression in PECs exposed to PAA compared to vehicle control treatment (Fig. 4a,f). We also showed no significant differences in expression of VCAM1 between PAA-exposed PECs and vehicle-treated SECs (Fig. 4a,f). Furthermore, PAA demonstrated reduced phosphorylation of eNOS^Ser1177^ (Fig. 4g), but a robust increase in eNOS^Thr495^ phosphorylation (Fig. 4h), in PECs following VCAM1 upregulation. To confirm the role of VCAM1 in the specific PAA regulation of eNOS signaling, we also tested the effect of VCAM1 silencing on PAA-exposed PECs. Knockdown of VCAM1 significantly increased the phosphorylation of eNOS at Ser1177, while decreased its phosphorylation at Thr495 in PECs treated with PAA (Supplementary Fig. 6d). These findings suggest that the enhanced expression of VCAM1 regulated by cytosolic localization of HDAC4 may involve the downregulation of eNOS signaling.

To better explain our findings about the effects of PAA on eNOS activation, which is thought to be regulated by elevated H_2_O_2_ levels, we next examined how DAAO-generated H_2_O_2_ in PECs (treated with D-alanine) modulate VCAM1-eNOS signaling. Our data support the notion that increased VCAM1 expression induced by chemogenetic generation of H_2_O_2_ in the cell mitochondria impairs nitric oxide (NO) signaling axis by eNOS^Ser1177^ dephosphorylation and increased phosphorylation of eNOS^Thr495^ following addition of D-alanine (Supplementary Fig. 5g-i). The similar outcome has been seen earlier in Fig. 4 in response to PAA treatment of PECs. In line with this, DAAO-generated H_2_O_2_ also led to a significant increase in the numbers of SA-β-gal^+^ cells (Supplementary Fig. 8a) and expression of γ-H2A.X (Supplementary Fig. 8b,c), indicating premature senescence and telomere shortening in ECs.

#### PAA Suppresses Angiogenic Capacity of ECs

Because EC senescence has been implicated in impaired angiogenesis, we reasoned that PAA might confer critical effects on angiogenic capacity of proliferating ECs. We, therefore, assessed intrinsic angiogenic incompetence in PECs exposed to PAA. The results show a markedly lower migration of PAA-exposed PECs (∼2-fold), as seen in SECs, compared to PECs 16 h post-scratch (Fig. 5a). We also tested the effects of PAA on tube formation ability, an in vitro measure of angiogenesis, of PECs. PAA exhibited a significant inhibition of tube formation, represented by a ∼5-fold decrease in the numbers of tubes formed by ECs, whereas PECs formed lower numbers of tubes (Fig. 5b). In agreement with this, aortic rings obtained from young mice which were incubated with PAA and exhibited markedly decreased endothelial sprouting compared to what seen in young aortic rings; while, no diLerences in numbers of endothelial sprouts were observed in old aortic rings (Fig. 5c). Sprouted aortic rings were also stained for endothelial-specific adhesion molecules VE-cadherin (VEC) and CD31, to better analyze the total volume of sprouted VEC- and CD31-expressing ECs. These data confirm that PAA reduces angiogenic capacity of young aortas by decreasing total volume of sprouted VEC^+^, CD31^+^ ECs (Supplementary Fig. 9).

**Fig. 5.**
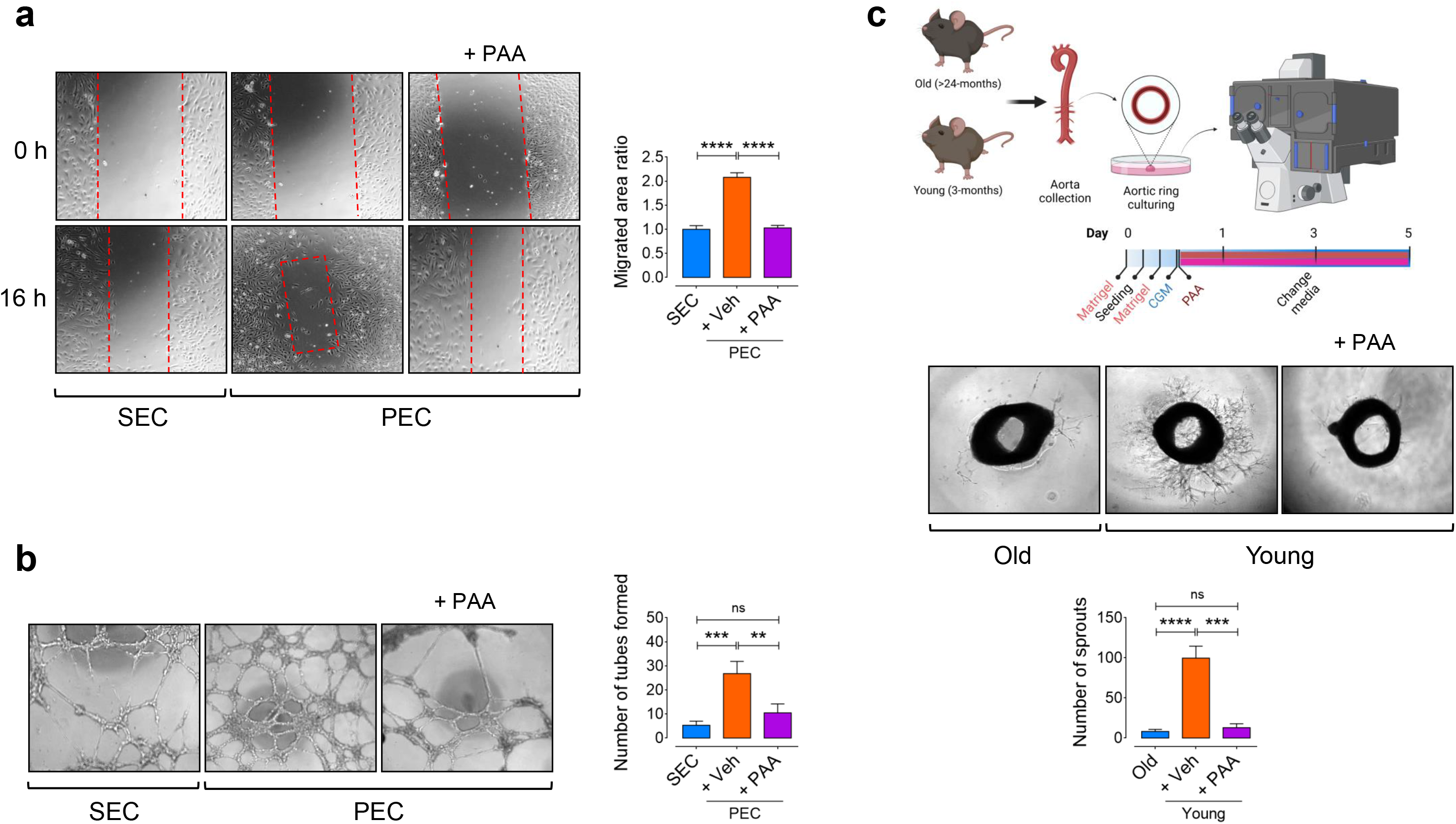
PAA reduces EC proliferation and angiogenesis. **a,** Confocal micrographs depict that PAA markedly reduces cell migration in PECs, shown as larger areas of uncovered surface, to the comparable level seen in replicative SECs. Right, Bar chart represents the ratio of cell migrated area (n=6). **b,** Confocal images represent the 2-D matrigel tube formation of PECs treated with vehicle or PAA and replicative SECs. Right, quantitative plot shows a significantly lower number of tubes formed by PAA-exposed PECs compared to vehicle-treated (n=6). **c,** Confocal micrographs of the aortic rings from old and young mice show that PAA treatment of young aortas significantly reduces angiogenic capacity compared to vehicle-treated young aortas. Bottom, quantitative plot is shown for the number of aortic endothelial sprouts (n=6). Scale bars, 200 μm. Data were determined in 6 micrographs from 3 different plates and represent triplicated biologically independent experiments. Error bars represent SD (**a-c**). *P* values were calculated using one-way ANOVA followed by Tukey’s post *hoc* test (**a-c**). (\*\**P*<0.01, \*\*\**P*<0.001, \*\*\*\**P*<0.0001). Source data are provided as a Source Data file.

#### The Preventive Effect of Acetate on PAA-Induced Premature Senescence in Endothelial Cells

To test how alteration of the gut microbiota-derived metabolites homeostasis upon aging underpins vascular endothelial senescence, we quantitated SCFAs including acetate. Corroborating our previous findings^18^, the concentrations of acetate in fecal samples obtained from old mice were markedly lower compared to what quantitated in young mice (Fig. 6a). We hypothesized that acetate might rescue senescence in endothelial cells exposed to PAA. Therefore, we explored a possible anti-senescence efficacy of acetate. HAEC were exposed to sodium acetate (3 μM, 3 days) upon treatment with PAA (10 μM) (Fig. 6b). As expected, acetate markedly reduced numbers of SA-β-gal^+^ cells in the presence of PAA (Fig. 6c) with senescence-related morphological transformations (Supplementary Fig. 3). In line with this, acetate strongly prevented PAA-induced telomere shortening by decreasing γ-H2A.X phosphorylation (Fig. 6d). Quantitative real-time PCR indicated that acetate effectively reduced mRNA levels of cell-cycle arrest markers p16^INK4a^ (Fig. 6e) and p21^WAF1/Cip1^ (Fig. 6f), and suppressed the SASP profile includingIL1β (Fig. 6g) and IL6 (Fig. 6h) triggered by PAA. Our data revealed that a decrease in IL6 expression (exposed to acetate) has been accompanied by reduced CaMKII^T286^ phosphorylation of HDAC4^S632^ and the expression of VCAM1 in PAA-treated PECs (Fig. 6i). VCAM1 immunostaining of acetate-treated PECs in the presence of PAA provided a proof of the concept that acetate-mediated HDAC4^S632^ dephosphorylation and its nuclear localization can boost the repressive effect of HDAC4 on VCAM1 and decrease its expression in these cells (Fig. 6j). These led to a significant increase in the phosphorylation of eNOS at Ser1177, while its phosphorylation at Thr495 was reduced (Fig. 6i).

**Fig. 6.**
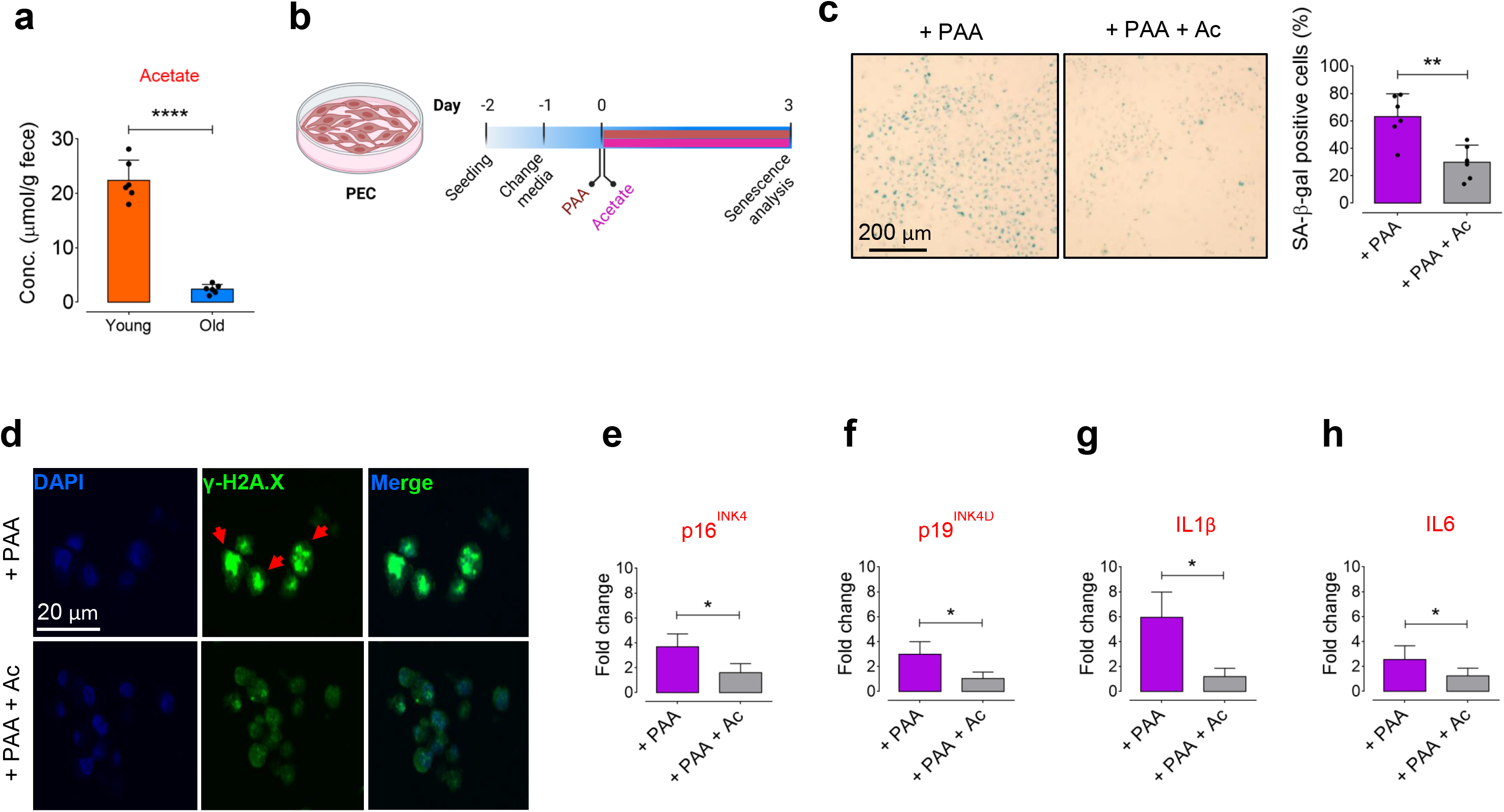
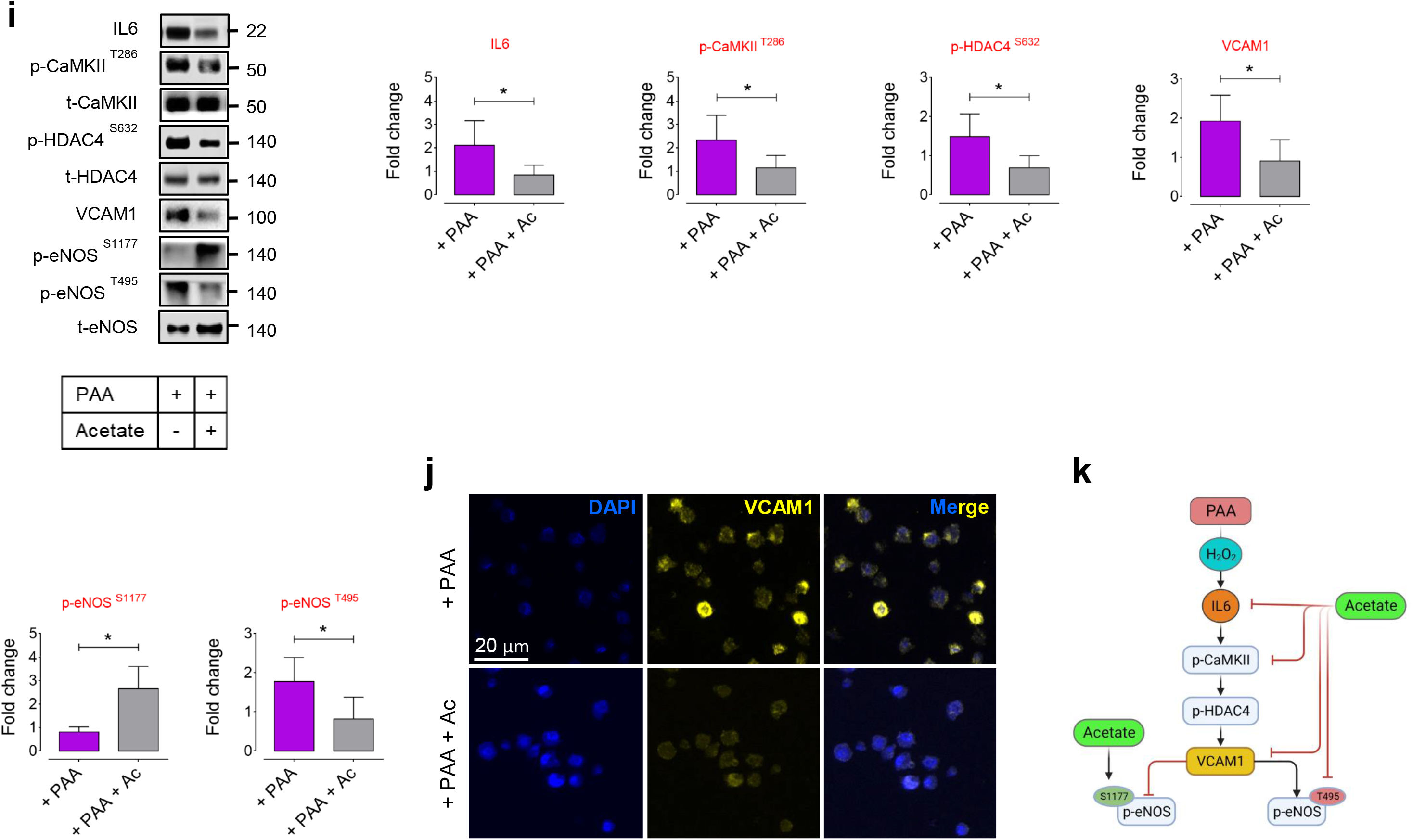
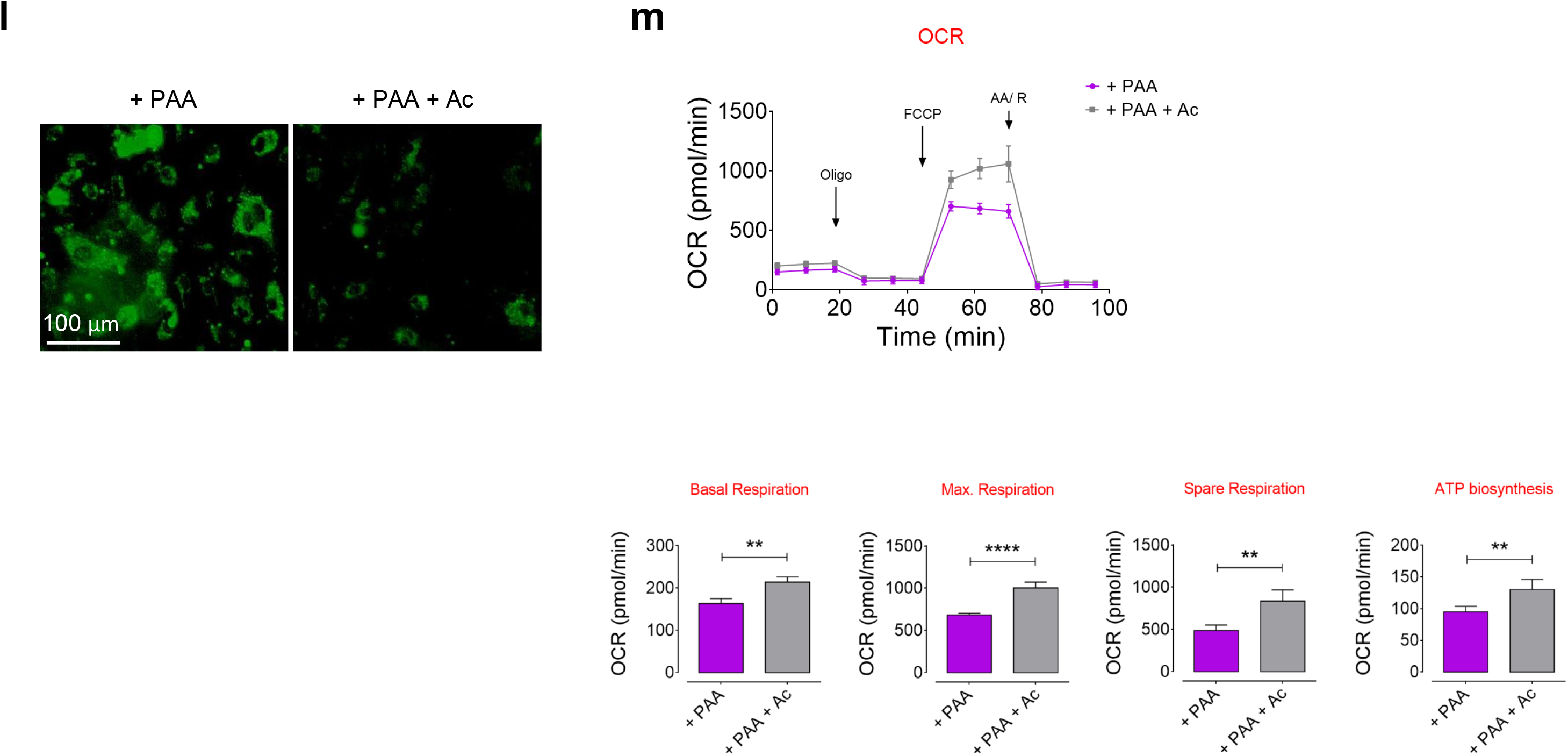
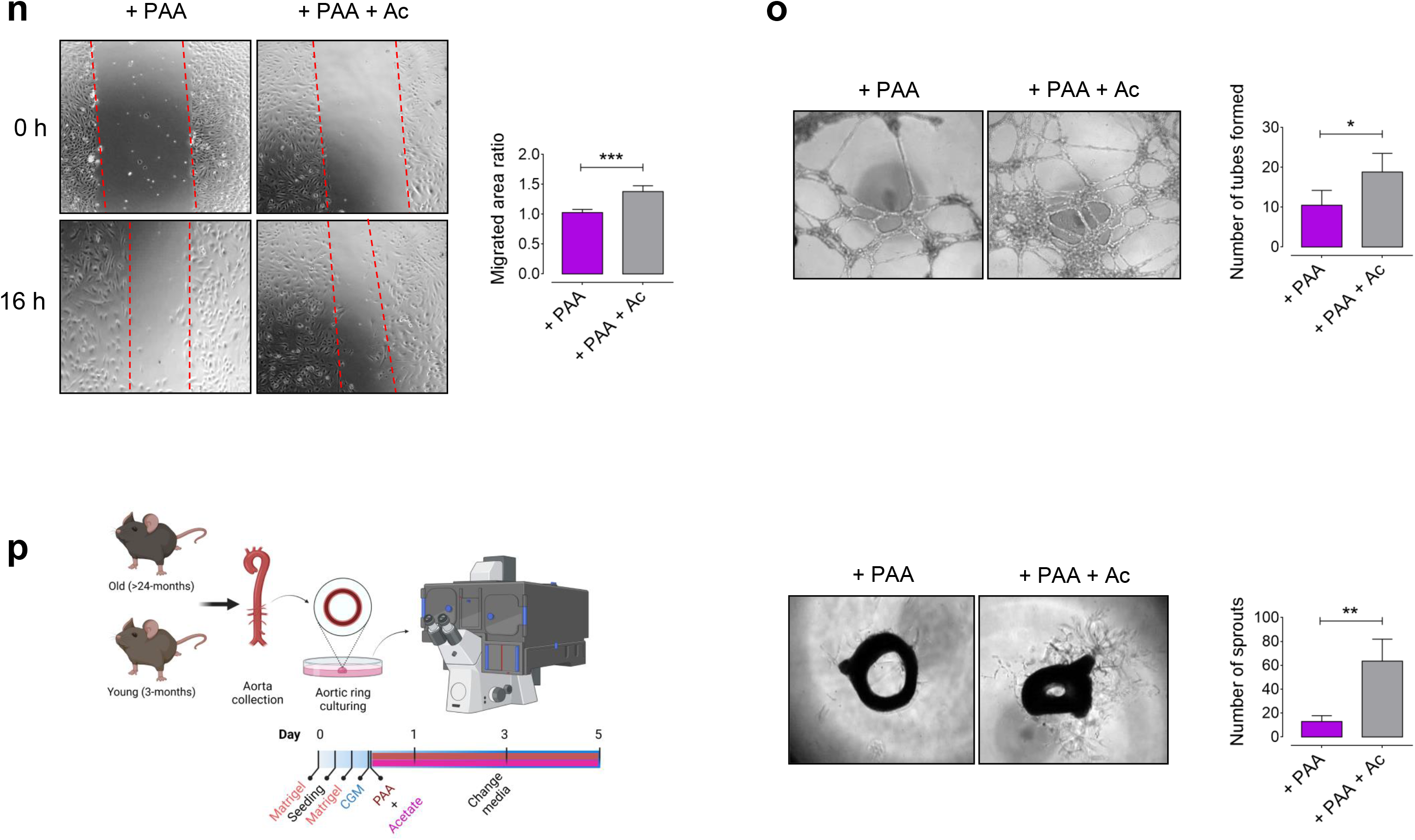
Acetate rescues PAA-induced EC senescence and restores angiogenic capacity. **a,** Fecal samples from old and young mice were collected for quantification of acetate by HPLC-RI. Bar chart represents significantly lower levels of acetate in feces of old mice compared to young mice (n=6). **b,** Schematic diagram of the experimental setting: PECs were co-treated with PAA (10 μM) and sodium acetate (3 μM) for 72 h and then analyzed for senescence hallmarks. **c,** Representative images of SA-β-gal staining (left) and bar chart (right) demonstrate that sodium acetate reduces cellular senescence in PAA-exposed PECs, represented by SA-β-gal-positive cells (%) (n=6). **d,** Representative images of γ-H2A.X foci demonstrate a marked reduction in DDR in PECs co-treated with PAA and sodium acetate compared to PECs exposed to PAA alone (n=6). **e-h,** qPCR analysis reveals that sodium acetate downregulates PAA-induced CDK inhibitors, p16^INK4a^ (**e**) and p19^INK4d^ (**f**), and SASP genes, IL-1β (**g**) and IL-6 (**h**), in PECs (n=6). **i,** Immunoblots show that sodium acetate regulates IL6-stimulated CaMKII-HDAC4 phosphorylation and the subsequent VCAM1 expression and eNOS phosphorylation at specific sites Ser1177 and Thr495 in PAA-exposed PECs (n=6). **j,** VCAM1 immunofluorescence represents that sodium acetate reduces its overexpression stimulated by PAA in PECs (n=6). **k,** Model for the proposed role of sodium acetate in restoration of eNOS signaling pathway in PAA-exposed PECs. We propose that acetate-mediated downregulation of IL6 is accompanied by reduced CaMKII-HDAC4 phosphorylation that localizes HDAC4 in the cell nucleus and subsequently dampens VCAM1 overexpression and restores eNOS signaling. **l,** CellRox green fluorescence staining demonstrates acetate-mediated reduction in intracellular ROS generation in PECs in the presence of PAA. **m,** Bioenergetics analysis by Seahorse metabolic analyzer reveals mitochondrial OCR in PAA-*versus* acetate+PAA-treated PECs. Bar charts show that co-incubation of PECs with sodium acetate and PAA restores mitochondrial function, represented by significant increases in basal and maximal OCR, spare reserve, and ATP biosynthesis as opposed to PAA-exposed PECs (n = 6). **n-p,** Confocal micrographs depict markedly positive effects of sodium acetate to restore endothelial angiogenic capcity, represented by cell migration (**n**), tube formation (**o**), and aortic endothelial sprouting (**p**). Quantitative plots represent significantly higher PEC migrated area ratio (n), the number of tubes formed (o), and the number of endothelial sprouts from aortic rings in the acetate+PAA-treated group compared to PAA-exposed group. Scale bars, 20, 100, and 200 μm. Error bars represent SD (**a,c,e-I,n-p**) or SEM (**m**). Data represent triplicated biologically independent experiments. *P* values were calculated using a two-tailed unpaired Student’s *t*-test. (\**P*<0.05, \*\**P*<0.01, \*\*\**P*<0.001, \*\*\*\**P*<0.0001). Source data are provided as a Source Data file.

We assessed the role of acetate in lowering ROS levels, particularly mitochondrial ROS, and improving mitochondrial respiration in PAA-induced senescent ECs, a finding of therapeutic relevance to counter EC dysfunction. The addition of acetate to PECs (exposed to PAA) prevented mitochondrial oxidative stress represented by lower ROS generated in the cell mitochondria (CellRox^+^ cells; Fig. 6l), whereas increased *Gpx1* transcripts to the level seen in vehicle-treated PECs (Supplementary Fig. 4). Accordingly, acetate improved mitochondrial dysfunction aggravated by PAA, shown as a roughly 30-40% increase in basal and maximal respiration rates and spare respiratory capacity. Moreover, ATP was biosynthesized at a significantly higher level in PAA-exposed ECs when supplemented with acetate (Fig. 6m). Notably, in proliferating ECs, acetate is essential for generating acetyl-coA to sustain the TCA cycle and redox homeostasis. Additionally, acetate might rescue the elevated ROS levels by stimulating fatty acid β-oxidation (FAO) in these cells^20^. Our data prove that acetate can counteract imbalanced oxidative stress in the cell mitochondria and restore its respiration capacity and energy supply in senescence ECs.

To further validate the positive effects of acetate on EC dysfunction, which was induced upon administration of PAA, we evaluated angiogenesis. Acetate promotes PAA-induced senescent ECs to become angiogenic. Our data show that acetate restores migration ability of PAA-exposed ECs (Fig. 6n) and improves *in vitro* angiogenesis represented by significantly increased numbers of tubes formed by senescent ECs induced by PAA (Fig. 6o). Moreover, in murine aortic rings, acetate induces endothelial sprouting responses *ex vivo*. Interestingly, 3-day treatment with acetate markedly increased numbers of sprouted VEC^+^, CD31^+^ ECs at day 6 in PAA-exposed young aortic rings on matrigel (Fig. 6p, Supplementary Fig. 9).

**Fig. 7.**
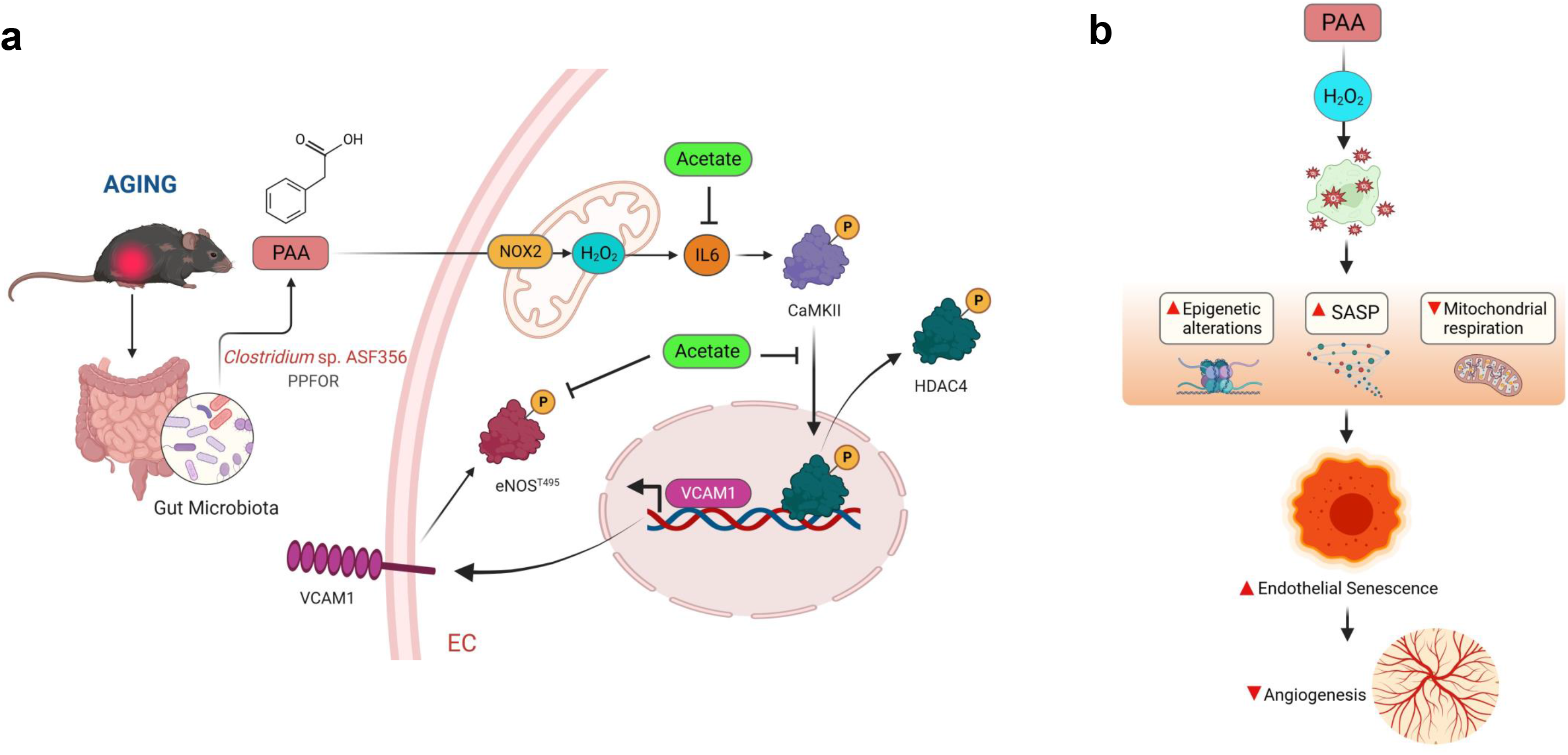
Proposed model for regulation of endothelial cell senescence by gut microbiota-derived PAA in aging. **a,** The mechanism underlying the contribution of gut-derived PAA to premature endothelial cell senescence. *Clostridium* sp. ASF356 harboring PPFOR gene homolog contributes to PAA generation in old mice. Age-dependently higher level of PAA regulates the SASP-epigenetic pathway which, in turn, augments cellular senescence. As an interventional strategy, acetate rescues PAA-induced premature senescence by regulating the SASP-mediated HDAC4 nuclear export and mitochondrial function, and thereby improves angiogenesis. **b,** PAA, through mitochondrial H_2_O_2_ production, modulates mitochondrial respiration and epigenetic regulation of the SASP, by which induces premature endothelial senescence and impairs angiogenic capacity in old organisms.

## Discussion

In this study, we examined the contribution of the gut microbiota and its derivative PAA to aortic endothelial senescence by combining shotgun metagenomics and targeted metabolomics analyses on plasma with cellular senescence and signal transduction studies in mice and human aging cohorts. Our combined metabolic and genetic characterizations reveal that the age-related increase in plasma levels of PAA and its derivative PAGln correlated with the presence of PPFOR/VOR gene homologs in the aged microbiome. These genes are responsible for catalyzing the oxidative decarboxylation of phenylpyruvate into a phenylacetyl-CoA intermediate, and ultimately into PAA (presumably by acyl-CoA thioesterase). Additionally, our findings revealed that age-related increase in circulating PAA levels was positively associated with higher abundance of *Clostridium* sp. ASF356 harboring PPFOR gene in aged microbiome. We screened the PAA-producing capacity of *Clostridium* sp. ASF356 *in vitro* as a proof of the concept and detected a remarkable generation of the metabolite in the presence of its precursor, phenylalanine. Our findings also confirmed that PAA induces morphological and metabolic changes in aortic ECs that lead to cellular senescence and consequently impairment of angiogenesis. Corroborating these results, our and others’ previous studies demonstrated that accumulation of senescent ECs plays a causative role in pathogenesis of vascular diseases such as atherosclerosis^1,33^. Therefore, the gut-derived PAA can be potentially suggested as a key determinant in the induction of endothelial senescence and the development of atherosclerosis upon aging.

PAA has long been recognized as a phenylalanine-derived precursor of PAGln in humans^34^ and phenylacetylglycine (PAGly) in rodents^22^, which is known as ’phenylketonuria metabolite’ associated with cognitive and neurodegenerative disorders^35^. In the context of cardiovascular diseases, PAA is primarily acknowledged for its role as a precursor in the generation of PAGln. Consequently, the *PPFOR* and *PPDC* genes have recently been linked to the prevalence of atherosclerotic CVD in patients exhibiting increased plasma PAGln concentrations^12^. However, only one human study has positively associated elevated plasma levels of PAA with the risk of incident major adverse cardiovascular events (MACE)^36^. Thus far, our study is the first to report the causal association between increased PAA with age and vascular senescence and dysfunction, as well as specific role of *Clostridium* sp. ASF356 in its formation in an aging model.

The present findings suggest novel mechanisms through which the clinical association between heightened PAA and vascular dysfunction in aging may occur. PAA confers its deleterious impact on proliferating ECs by aggravated oxidative stress as well as the expression of the CDK inhibitors p16^INK4a^, p19^INK4d^, and p21^WAF1/Cip1^, known regulators of cellular senescence^4^, which was also seen in aortic CD31^+^-endothelium of old mice. By excessive ROS production, particularly H_2_O_2_, in the mitochondria of PECs, PAA induces the expression of these CDK inhibitors and increases SA-β-galactosidase activity and DNA damage response (represented by increased γ-H2A.X), and thereby promotes EC senescence. Aside from this, PAA-induced stress condition counteracts ROS-scavenging system including GPx1, which converts oxidized glutathione (GSSG) to its reduced form (GSH), in PECs^24^. These data are consistent with what observed in oxidized low-density lipoprotein (oxLDL)-induced senescent HUVEC endothelial cells^37^, in which glutathione peroxidase decline has been associated with a dramatic activation of p53/p21 pathway and subsequent endothelial dysfunction, proven as subclinical stage of atherosclerosis^38^. Thus, we believe that PAA may potentially be a pro-oxidant inducer of EC senescence in aging organisms. Beyond oxidative stress, senescent cells develop another senescence hallmark which is a persistent proinflammatory phenotype that impairs cellular function^5^. Our *in vitro* studies strongly suggest that excess mitochondrial H_2_O_2_ triggers pro-inflammatory cytokines (IL1α, IL1β, and IL6) and cell adhesion molecule VCAM1 in ECs following treatment with PAA. To confirm that PAA-induced senescence-messaging secretome is H_2_O_2_-dependent, we exploited a chemogenetic approach utilizing a recombinant lentiviral DAAO to generate H_2_O_2_ in the endothelial cell mitochondria. We observed an increased expression of IL6 in response to chemogenetic H_2_O_2_, explaining that PAA upregulates SASP through a H_2_O_2_-dependent fashion. However, the mechanisms which underlie the H_2_O_2_-mediated SASP in PECs exposed to PAA are still quite a challenge to uncover.

To explore potential answer for this question, we hypothesized that PAA, by upregulating mitochondrial H_2_O_2_-IL6, regulates epigenetic-SASP responses in ECs. Therefore, we examined HDAC4-mediated epigenetic regulation of VCAM1-eNOS pathway in PAA-exposed ECs. Our findings revealed that PAA aggravates phosphorylation of HDAC4 and its nuclear export towards the cell cytosol. In accordance to prior studies, NADPH oxidase 4 (NOX4)-produced H_2_O_2_ regulates HDAC4 shuttle between cell cytosol and nucleus through two potential mechanism, one direct oxidation of HDAC4, and another one H_2_O_2_-regulated phosphorylation of HDAC4^30^. Matsushima and colleagues discovered that phosphorylation of HDAC4, secondary to its oxidation, facilitates nuclear export^39^. Beyond, we observed that protein kinase CaMKII regulates HDAC4 phosphorylation at Serine site 632, which subsequently controls its cytosolic accumulation following treatment with PAA. In line with this, protein kinases such as CaMKII family have been demonstrated to stimulate multisite (including Ser632) hierarchical phosphorylation of HDAC4 and to facilitate its subcellular localization. Furthermore, PAA-exposed senescent ECs exhibit that IL6 upregulation is associated with increased CaMKII phosphorylation. Consistent with this observation, treatment with IL6 and its soluble receptor (sIL6R) also resulted in a significant increase in the phosphorylation of CaMKII in a JAK/STAT3-dependent fashion in HUVEC that regulates endothelial motility and proliferation^40^. Therefore, we believe that PAA both directly, by oxidizing, and indirectly, by regulating CaMKII, phosphorylates HDAC4, and thereby regulates de-repression of its downstream adhesion molecule VCAM1 in ECs. Our *in vitro* experiments showed that PAA-mediated HDAC4 nuclear export disrupts HDAC4/VCAM1 complex, through which increases *Vcam1* transcription. These data support the hypothesis that the effect of PAA on EC dysfunction is probably mediated through VCAM1 upregulation. Unfortunately, whether HDAC4 nuclear export regulates *Vcam1* transcription was not well understood, although previous report by Yang and colleagues (2018) demonstrated that genetic downregulation of HDAC4 leads to reduced VCAM1 protein expression in ECs following treatment with angiotensin-II^32^. We finally showed that the epigenetic regulation of VCAM1 promotes differential phosphorylation responses for specific phosphorylation sites (Ser1177 and Thr495) on eNOS, which have been involved in EC senescence and dysfunction. As reported previously, genetic deletion of eNOS disrupts VEGF-induced angiogenesis *in vivo*, whereas phosphorylation of eNOS at Ser1177, but not Thr495, by multiple protein kinases increases neovascularization in ischemic tissues^41^. We, therefore, suggest that higher PAA levels during aging negatively impact angiogenesis likely through reducing eNOS function, leading to age-related endothelial dysfunction. Our results demonstrate that PAA actively impairs angiogenesis represented by a markedly reduction in EC migration and tube formation *in vitro*, as well as a decrease in aortic endothelial sprouting *ex vivo*, accompanied by increased EC senescence phenotype. Angiogenesis assumes a heightened significance, particularly when the process becomes compromised in older patients with arterial stiffness, atherosclerosis, or ischemic heart disease^42^. In these clinical scenarios, impaired angiogenesis fails to elicit the necessary compensatory mechanisms required for augmenting blood supply in conditions characterized by reduced blood flow and compromised tissue perfusion. This deficiency underscores the urgency of understanding and harnessing angiogenesis as a successful treatment of aging-related cardiovascular pathologies.

Since oxidative stress alters pathways sensing changes in cellular energy balance in dysfunctional and senescent ECs^43^, we next determined whether mitochondrial dysfunction induced by PAA, as shown to promote excessive H_2_O_2_ in mitochondria, can explain the negative effects of PAA on angiogenic capacity of ECs. When proliferating endothelial cells become angiogenic, they upregulate fatty acid β-oxidation to sustain the TCA cycle and mitochondrial respiration for energy supply^20^. Moreover, endothelial cells rely on glycolysis for energy production^26^ and increase it to promote cell migration and proliferation^44^. In contrast, pathways promoting permanent proliferative arrest, including oxidative stress, have been demonstrated to shut down both OXPHOS and glycolysis and to impair ATP biosynthesis in senescent endothelial cells^45^. Our results revealed that PAA exposure of PECs reduces their mitochondrial respiration, as detected by OCR, and glycolysis, as detected by ECAR. Hence, this might confirm that PAA-induced senescent ECs would exhibit mitochondrial metabolic alterations, so that a strategy to induce mitochondrial reprogramming in the cells may potentially restore senescence-related impaired angiogenesis.

Here, we characterized how acetate can reprogram the PAA-treated PECs’ metabolism when switching from proliferation to senescence. Amidst the spotlight on other gut-derived SCFAs, acetate emerges as an influential molecule in the landscape of aging-related cardiovascular diseases. A research by Kalucka et al. (2018) has unveiled acetate’s role in rescue of fatty acid β-oxidation in quiescent ECs and restoring endothelial function^20^. Moreover, acetate supplementation has clinically been associated with reduced arterial stiffness^46^ and lower blood pressure^47^ in aged mice. Mechanistically, acetate (converted to acetyl-coA) might counteract ROS-induced oxidative stress and rescue mitochondrial dysfunction^20^. Acetate supplementation can be considered as a microbiome-based postbiotic strategy for reversal of gut-derived PAA-induced EC senescence: It is simply utilizing the gut microbiome for alleviating gut microbiome-related vascular dysfunction in an old host. Accordingly, we tested whether sodium acetate can rescue PAA-induced EC senescence and restore angiogenic capacity. Our data suggest that exogenous acetate supplementation of PAA-exposed PECs rescued cellular senescence, lowered H_2_O_2_-induced oxidative stress and restored mitochondrial respiration to maintain a high energetic proliferating state in PAA-exposed ECs. Because of an association between shortened telomeres and impaired mitochondrial energy supply in aged ECs^48^, we also demonstrated that acetate reduces DNA damage (reduced γ-H2A.X phosphorylation), which is probably mediated by downregulating CDK inhibitors p16^INK4a^ and p19^INK4d^. Consistently, the p16^INK4a^-expressing cells have been shown to accumulate in the vasculature and heart, and negatively influence lifespan in INK-ATTAC mice^49^. In contrast, elimination of p16-positive senescent ECs reduces DNA damage, delays atherosclerosis, and notably extends the median lifespan in progeroid mice^50^. We further observed that acetate preserves the profound reductions observed in basal and maximal oxygen consumption rate in the presence of PAA, suggesting that acetate reinforces these cells to compensate for decreased ATP supply via increased mitochondrial respiration to maintain cellular energy homeostasis essential for angiogenesis. Acetate could also regulate epigenetic-SASP interplay in PAA-treated ECs, where it downregulates VCAM1 and alleviates endothelial dysfunction. As we and others exhibited, IL6 underlies vascular premature aging and atherosclerotic CVD^24^, whereas acetate at least in HUVEC can inhibit the production of IL-6 and IL-8 and subsequently reduce VCAM1-mediated cell adhesion^51^. In line with this, our findings revealed that acetate, by downregulating IL6-CaMKII, reduces HDAC4^S632^ phosphorylation and stabilizes its nuclear localization, which thereby decreases VCAM1 expression. Indeed, the outcome was an increase in eNOS phosphorylation at Ser1177, which has long been recognized to regulate endothelial sprouting and neovascularization *in vivo* through enrichment of genes related to cell polarity such as Partitioning Defective (PARD) family detected at the single-cell level^42^. eNOS has also been shown to regulate angiogenesis in a hind limb ischemia model, in which the endothelial-specific knockout of Rac1, a Rho family member, reduces eNOS activity and NO generation that finally abolishes aortic capillary sprouting and peripheral neovascularization^52^. Therefore, in addition to restoration of mitochondrial respiration and energy supply, acetate may restore angiogenic capacity of ECs likely through suppressing VCAM1 and increasing eNOS function. Our findings suggest acetate as an effective senotherapeutic compound with potential pro-angiogenic efficiency for recovery of ischemic vascular diseases such as coronary artery disease and peripheral arterial disease in older adults.

Taken together, our findings reveal that the gut microbiota-derived PAA is a key determinant of EC senescence and the development of angiogenic decline in aging through a mitochondrial H_2_O_2_-regulated SASP and epigenetic modifications and subsequent mitochondrial dysfunction. We also suggest *Clostridium* sp. ASF356 harboring PPFOR homologs to regulate circulating PAA in aged mice. Importantly, we propose that the bacteria and its byproduct PAA hold an independent predictive value for cardiovascular risk. A personalized bacterial genetic engineering for disruption of PPFOR may suggest a novel therapeutic strategy for aging-related prevention of endothelial dysfunction and age-related CVD in the elderly. However, these approaches require further development.

## Methods

### Mice

Our research complies with all ethical regulations approved by the Institutional Animal Care and Use Committee at University of Zurich. Animal care and all experimental protocols were in accordance to the Directive 2010/63/EU of the European Parliament and of the Council of 22 September 2010 on the protection of animals used for scientific purposes.

8 week old wild-type C57BL/6J female and male mice were purchased from the Jackson Laboratory and maintained in our facilities for specific times for normal aging modeling. All mice were individually housed in controlled environments in plexiglass cages under a strict 12 h:12 h light/dark cycles at an ambient temperature of 23 ± 1°C and humidity of 55 ± 10%, and fed standard chow diet (including 19% Protein, 61% Carbohydrate and 7% Fat, #D11112201, Research Diets, New Brunswick, NJ, USA) until 3 (as Young) or >24 (as Old) months of age. Mice had access to drinking water and food *ad libitum*. The animals were monitored for body weight during their lifespan. The investigators in this study worked in a double-blinded manner.

### Blood and fecal collection, and tissue harvesting

Mice feces were collected and quickly stored at -80°C for DNA extraction and shotgun metagenomics and quantification of acetate levels.

Mice were euthanized by CO_2_ inhalation in an area separate from the housing facility. Whole blood was collected by cardiac puncture from mice. Plasma was isolated by centrifuging at 125 *g* for 8 min at room temperature, as previously described^18^. Plasma samples were used for LC-MS/MS quantification of PAA and PAGln. For measurement of serum creatinine and blood urea nitrogen (BUN), an aliquot of blood samples was incubated for 1 h at room temperature, and then centrifuged at 1500L×Lg for 10Lmin.

After blood collection, heart was perfused with normal saline and aorta was harvested, followed by removal of adhering connective and fat tissues. Aortas were kept in Opti-MEM + 2.5% (v/v) fetal bovine serum (FBS) for *ex vivo* vasorelaxation and aortic endothelial sprouting. For immunohistochemical and signaling analyses, aortic tissues were quickly fixed in 10% formalin or snap-frozen in liquid nitrogen followed by storage at -80 LC, respectively.

### Mice creatinine and BUN measurement

Serum creatinine was measured using a colorimetric detection kit (Cayman Chemical; 700460). Serum samples were treated with creatinine reaction buffer (including Creatinine Sodium Borate, Creatinine surfactant, and Creatinine alkaline buffer), followed by a color reagent, according to the manufacturer’s instructions. Absorbance was read at 495 nm with a microplate reader.

Moreover, sera were diluted and incubated with specific color reagents for colorimetric BUN measurement, according to the manufacturer’s instructions (Invitrogen; EIABUN), and absorbance was read at 450 nm.

### Human metabolomic data (TwinsUK Cohort)

To validate the associations between circulating PAA and PAGln concentrations and age, we used the metabolomic data of TwinsUK human aging cohort^21^. The TwinsUK cohort encompasses metabolome data of 1,116 metabolites (including PAA and PAGln) profiled in more than 14,000 participants between 18 and 95 years old. Participants that exhibited values below the detection level (0) for PAA or PAGln were considered as not available. Accordingly, PAA and PAGln were measured in plasma samples of 7,303 subjects by Metabolon using a non-targeted UPLC–MS/MS platform.

### Targeted LC–MS/MS analysis of plasma PAA and PAGln levels

PAA and PAGln concentrations in mouse plasma (100 μL) were quantitated by liquid chromatography coupled to a high resolution mass spectrometer (LC-HRMS). Briefly, frozen ethanol/methanol (-20°C) containing internal standards (D5-phenylacetic acid and D5-phenylacetylglutamine at concentration of 0.25µg/ml) was added to plasma samples, followed by centrifugation at 13,000 rpm at 4°C for 20 min. Supernatant was dried under nitrogen at 30°C and reconstituted into 100µl of water/methanol (90/10) for next centrifugation at 13,000 rpm for 20 min. Supernatants were separated in a Waters HSST3 column (150 mm × 2.1 mm; 1.8 μm) for LC-MS/MS analysis (Thermo Ultimate 3000 system hyphenated with a Thermo Exploris 120 orbitrap mass spectrometer) using a solvent gradient of buffer B (Ammonium acetate 1 mM in acetonitrile) to buffer A (Ammonium acetate 1 mM in water). The injected volume was 5µl and the calibration range was 10 to 2000 ng/ml. Peaks were identified based on standards for each of the metabolites and processed using Tracefinder software. All MS conditions are shown in Table 1 and were optimized with the use of standard solutions.

**Table 1.** Plasma PAA and PAGln mass spectrometry conditions.

|  | Polarity | Formula | Adduct | Precursor ion (m/z) | Product ion (m/z) | HCD (%) |
| --- | --- | --- | --- | --- | --- | --- |
| PAA | Negative | C<sub>8</sub>H<sub>8</sub>O<sub>2</sub> | -H | 135.0452 | 91.0553 | 30 |
| PAA-d5 | Negative | C<sub>8</sub>H<sub>3</sub>O<sub>2</sub>D<sub>5</sub> | -H | 140.0765 | 96.0869 | 30 |
| PAGln | Negative | C<sub>13</sub>H<sub>16</sub>N<sub>2</sub>O<sub>4</sub> | -H | 263.1037 | 127.0513 | 60 |
| PAGln-d5 | Negative | C<sub>13</sub>H<sub>11</sub>N<sub>2</sub>O<sub>4</sub>D<sub>5</sub> | -H | 268.1351 | 127.0513 | 60 |

### Fecal DNA extraction

Fecal genomic DNA was extracted as described previously^18^. 122 μL MT buffer (MP Biomedicals) in 978 μL was added to 200 mg of fecal pellets in a sterile microcentrifuge tube, mix thoroughly and homogenized in FastPrep. The mixture was centrifuged at 14,000 ×Lg for 10 min. Then, 250 μL supernatant was transferred to a new sterile tube which containing equal volume of PPS buffer (MP Biomedicals), and centrifuged at 14,000 ×Lg for 5 min. A total of 730 μL liquid was aspirated onto a DNA purification column and centrifuged at 14,000 ×Lg for 1 min. After washing with SEWS-M washing buffer twice, the purified DNA was dissolved in 100 μL sterile Milli-Q water. The DNA concentration and purity were evaluated by a NanoDrop 2000 (Thermo Scientific).

### Shotgun metagenomics

Metagenomic DNA libraries were generated using a high-quality genomic DNA and libraries were sequenced on the Illumina NovaSeq 6000 PE150 platform with a 250-bp paired-end model. Briefly, samples were subjected to quality control by using Qubit^®^ dsDNA Assay Kit in Qubit^®^ 2.0 Flurometer (Life Technologies, USA). 1 μg of DNA samples were fragmented to 350 bp by sonication, end-polished, A-tailed, and ligated with the full-length adaptor for Illumina sequencing, followed by PCR amplification. DNA libraries were analyzed for integrity and purity using the Agilent2100 Bioanalyzer (Agilent, USA) and quantified via real-time PCR. The libraries were sequenced on an Illumina NovaSeq platform (Novogene, UK) and 250 bp paired-end reads were generated. llumina TruSeq adapters and low quality reads were trimmed using Trimmomatic (v0.39, parameters: *TruSeq3-PE-2.fa:2:30:10 SLIDINGWINDOW:5:20 LEADING:3 MINLEN:60*)^53^. Quality filtered reads were then aligned against the mice reference genome (https://ftp.ncbi.nlm.nih.gov/genomes/all/GCF/000/001/635/GCF_000001635.27_GRCm39/GCF_000001635.27_GRCm39_genomic.fna.gz) to remove host reads using Bowtie2 (*parameters: --very-sensitive-local*) and read pairs concordantly mapping the host genome were discarded.

#### Taxonomic profiling

Taxonomic profiling was performed using kraken2/Bracken2 (parameters: *--confidence 0.51*)^54^ against the mouse microbiota genome catalog (CMMG) (https://ezmeta.unige.ch/CMMG/Kraken2db/cmmg)^55^. Kraken2 profiles were then subjected to Bracken2 for taxonomic refinement using the corresponding database and read length. Bracken2 corrected CMMG read counts were then normalized for reference genome length and loaded together with available reference genomes taxonomy (https://ezmeta.unige.ch/CMMG/Supplementary_Tables_and_Figures/Tables/Table_S4_curated_taxonomy.tsv) into a phyloseq object for data handling. Only the taxa identified in at least 2 samples with coverage of at least 100 reads in total were considered for further analyses.

#### Functional profiling

Genomes were screened for presence of K00169 (*vor*) and or K00179 (*ppfor*), the genes encoding the enzymes participated in the phenylalanine metabolism and production of PAA, based on their available Kyoto Encyclopedia of Genes and Genomes (KEGG) functional annotations (https://ezmeta.unige.ch/CMMG/functional_annotations/mouse/annotations.tsv). The differential abundance of K00169 or K00179 in the identified genome was then added as additional information in the phyloseq object to identify *vor-* and/or *ppfor*-positive taxa in the microbiome of old and young mice. The association between the abundance of KO and plasma levels of PAA and PAGln were also characterized by a Spearman correlation analysis.

#### Community analysis

Microbiota profiles were characterized using alpha-diversity and beta-diversity^18,56^. Hill-d0, 1, and 2 indices represent the richness of detected taxa in the aged or young microbiome. Moreover, Pielou’s index (evenness) represents the relative abundance of each taxon in the communities. The influence of age and sex (=cage) was tested on alpha-diversity and beta-diversity metrics using Wilcox univariate nonparametric test and permutational multivariate analysis of variance (PERMANOVA) multivariate test. Age- and sex (=cage)-associated differences in taxa abundance were assessed at all taxonomic ranks using analysis of composition of microbiomes (ANCOM).

#### Fecal acetate quantification

Mice fecal pellets were stored at -80°C for quantification of acetate, using high performance liquid chromatography equipped with a refractive index detector (HPLC-RI)^18^. Briefly, feces were mixed with H_2_SO_4_ 10 mM, homogenized, and centrifuged at 6000 rpm for 20 min at 4°C. The supernatant was filtered and then separated with a LaChrom HPLC-System (Merck-Hitachi, Japan) using a SecurityGuard Cartridges Carbo-H (4 x 3.0 mm) (Phenomenex Inc., Torrance CA, USA) connected to a Rezex ROA-Organic Acid H^+^ column (300 x 7.8 mm; Phenomenex Inc.). The elution of samples (40 μL injection) was carried out at 40 °C under isocratic conditions (10 mM H_2_SO_4_, flow rate 0.4 mL/min), and analytes were quantified using a refractive index detector L-2490 (Merck Hitachi). EZChrom software was used for data processing.

### *In vitro* screening of PAA and PAGln formation by *Clostridium* sp. ASF356

Axenic culture of *Clostridium* sp. ASF356, a strain from the Altered Schaedler flora (ASF), was obtained from Prof. Michael Wannemuehler at the Iowa State University. *Clostridium* sp. ASF 356 was reactivated from -80°C glycerol stocks by a 1 % (v/v) inoculation in either Brain Heart Infusion (BHI) or Yeast Casitone Fatty Acids (YCFA) media anaerobically at 37°C. Overnight saturated pre-cultures were transferred (1%, v/v) into the fresh media, and cells were grown for 46 hours at 37°C. Bacterial cultures were then centrifuged (12,000 × g for 10 min at 4°C), and the supernatant was collected and stored at -80°C for further LC-MS/MS quantification of PAA and PAGln. Standard-containing supernatants were centrifuged at 13,000 rpm at 4°C for 20 min, and then separated in a Waters HSST3 column (150 mm × 2.1 mm; 1.8 μm) for LC-MS/MS analysis (Thermo Ultimate 3000 system hyphenated with a Thermo Exploris 120 orbitrap mass spectrometer; volume of injection: 5 μl) using a solvent gradient of buffer B (Ammonium acetate 1 mM in acetonitrile) to buffer A (Ammonium acetate 1 mM in water). Peaks were characterized and analyzed based on internal standards (Table 1) for each of the metabolites and processed using Tracefinder software.

### Cell Line

The primary human aortic endothelial cells (HAEC) were grown in EBM-2 endothelial cell growth basal medium supplemented with EGM-2 cell growth supplement pack containing FBS (10% v/v), L-glutamine (2 mM) and penicillin–streptomycin (100 μg/mL) and incubated at 37°C in 5% CO_2_. The cell line was tested to exclude any positive status for mycoplasma. Cells were studied in passages 4-5 (proliferating) and 15-17 (replicative senescence). The cellular senescence was verified by SA-β-galactosidase staining and immunoblots of p21^WAF1/Cip1^.

### Endothelial Cell Senescence Models

#### Replicative senescence

HAEC (p.4 or p.5) were seeded in 100% EGM-2 at a density of 5,000 cells/cm^2^ and the culture medium was changed every 48 h. To generate the corresponding proliferative control (PEC), cells were trypsinized and cultured in a growth medium for 48 h to proliferate. In parallel, we continued cell passage every 48 h until the proliferation is suppressed and replicative senescence phenotype (due to repetitive passages) is proved (SEC; p.15 to p.17). The cellular senescence was verified by SA-β-galactosidase staining.

#### H_2_O_2_-induced premature senescence

In order to induce premature senescence, proliferating HAEC (p.4 or p.5) were cultured in 100% EGM-2 with 20% fetal bovine serum (FBS) and treated with exogenous H_2_O_2_ (50 μM). After 4 h, the culture medium was changed, and cells were kept on culture for up to 72 h.

### SA-β-galactosidase staining

Cultured HAEC cells were stained for SA-β-galactosidase according to the manufacturer’s protocol (Merck Millipore). Briefly, cells were fixed for 15 min and then stained with SA-β-gal detection solution overnight at 37°C. Senescent cells, illuminated as blue-stained cells, were captured under a light Olympus microscope and analyzed using ImageJ software.

### CellRox green staining

Cells were grown to confluence in growth medium and treated with CellROX^®^ Green reagent at a final concentration of 5μM (Invitrogen, C10444) for 30 min at 37 °C to detect intracellular ROS by a Leica TCS-SP8 fluorescence microscope. The fluorescence intensity was evaluated using ImageJ software.

### Lentiviral DAAO-HyPer7.2 transduction

The HAECs were transduced with lentivirus-DAAO-HyPer7.2 targeted to the cell mitochondria at a MOI of 20-50 in serum-free culture media which was exchanged for fresh media containing 10% FBS after 5 hours. Cells were treated with D-alanine (10 mM) for the generation of mitochondrial H_2_O_2_ 48-72 hours after lentiviral transduction.

### HyPer7.2 fluorescence imaging

The cells expressing adenovirus serotype 5 (AV5)-HyPer7.2 targeted to the cell mitochondria at a MOI of 1000 were treated with specific concentrations of either exogenous H_2_O_2_ or PAA. Cells were then washed and incubated in a HEPES-buffered solution containing 140 mM NaCl, 5 mM KCl, 2LmM CaCl_2_, 1LmM MgCl_2_, 10 mM D-glucose and 1 mM HEPES (pH 7.4) for 2 hours at 37 °C. Coverslips were mounted on a live-cell imaging platform that allowed for stable superfusion.

For real-time fluorescence imaging, the ratiometric HyPer7.2 biosensor was excited at 420 nm and 490 nm, and emission was recorded at 530 nm using Metafluor Software, as previously described^57-59^. The mitochondrial H_2_O_2_ measurement was performed with a 20X oil immersion objective. Following background subtraction, mitochondrial H_2_O_2_ was defined as R/R_0_, ratio, where R is the ratio of the 490 nm to the 420 nm signals and R_0_ is the baseline 490/420 ratio.

### Real-time quantitative PCR

Total RNA from cells and aortas was extracted using TRIzol Reagent^®^ (Sigma-Aldrich), according to the standard protocol previously described^60^. cDNA was synthesized using High-Capacity cDNA Reverse Transcription Kit (Thermo Fisher Scientific) and amplified in a StepOnePlus RT-PCR thermocycler (Applied Biosciences) with Power SYBR Green PCR Master Mix (Thermo Fisher Scientific). Genes of interest were amplified with the corresponding gene-specific pairs of primers designed according to the coding strand of genes in the National Center for Biotechnology Information (NCBI) database using Integrated DNA Technologies (idtdna) and listed in Supplementary Table 1.

### siRNA transfection

HAECs were grown to 70–80% confluence for transfection with OnTARGETplus short-interfering RNA (siRNA) smart pool (Dharmacon) targeting CaMKII (siCaMKII, L-004942-00-0005) and VCAM1 (siVCAM1, L-013351-00-0005). OnTARGETplus Non-targeting scrambled control siRNA (siNeg, D-001810-10-05) was utilized for as negative control. Cells were transfected with siRNA using Lipofectamine RNAiMAX transfection reagent (Invitrogen) according to the manufacturer’s protocols. After 48 h, HAECs were collected for subsequent analysis. The silencing efficiency was confirmed by immunoblots.

### Antibodies

Antibodies against phospho-CaMKII Thr286 (1:1000; 12716), total-CaMKII (1: 1000; 4436), phospho-HDAC4 Ser632 (1:1000; 3424), total-HDAC4 (1:1000; 7628), H3 (1:1000; 9715), phospho-eNOS Ser1177 (1:1000; 9517), phospho-eNOS Thr495 (1:1000; 9574), total-eNOS (1: 1000; 32027), and phospho-Histone H2A.X Ser139 (1:1000; 80312) were purchased from Cell Signaling Technology; antibodies against VCAM-1 (1:1000; MA5-31965), IL6 (1:1000; M-620), NADPH oxidase 2 (NOX2) (1:1000; PA5-76034), and VE Cadherin (1:100; PA5-19612) were obtained from Invitrogen; antibody against CD31 (1:100; 14-0311-82) was purchased from eBioscience; antibody against Histone H3ac (Pan-Acetyl) (1:500; sc-518011) was obtained from Santa Cruz Biotechnology; antibody against p16^INK4A^ (1:100; ZRB1437) was purchased from Merck Millipore; antibody against phospho-HDAC4 Ser632 (1:100; ab39408-1001) and conjugated secondary antibodies against Alexa Fluor^®^ 488 (1:2000; ab150157) and Alexa Fluor^®^ 647 (1:2000; ab150075) were obtained from abcam; antibodies against goat anti-rabbit IgG-HRP (1:2000; 4030-05) and goat anti-mouse IgG-HRP (1:2000; 1036-05) were purchased from Southern Biotechnology.

### Immunoblotting

Proteins were extracted from in ice-cold RIPA lysis buffer (ThermoFisher Scientific) or nuclear and cytoplasmic extraction reagents (for protein isolation from nuclear and cytosolic fractions) supplemented with protease and phosphatase inhibitors cocktails (ThermoFisher Scientific). Equal amounts of protein lysates (20 µg) were separated in 10% SDS-PAGE gels and transferred to PVDF membranes (Bio-Rad). The membranes were incubated with specific primary antibodies (1:1000) overnight at 4°C and then with corresponding horseradish peroxidase (HRP)-labeled secondary antibodies (1:2000) for 1h. The protein bands were visualized using an Immobilon Western HRP substrate Crescendo and Forte Reagents and Amersham Imager 600 (GE Healthcare). Quantitative densitometric analyses were performed using ImageJ software.

### Opal multiplex immunofluorescence

Murine aortas and aortic endothelial cells were formalin-fixed and paraffin-embedded. Experimental sections (4-μm thickness) and positive tissues were stained with specific primary antibodies (1:100) followed by Opal multiplex immunostaining system (Akoya Biosciences) to generate the stained slides. The staining conditions for all primary antibodies were optimized using chromogenic DAB detection (Leica Biosystems). Using an Opal 7-Color Automation IHC Kit (Akoya Biosciences), Opal singleplex and multiplex for initially-optimized antibodies p-HDAC4^Ser632^, CD31, p16^INK4A^, VCAM1, and γ-H2A.X^Ser139^, as well as DAPI (for nuclear counterstaining) were conducted on a Leica Bond Rx automated autostainer (Leica Biosystems). The panel of primary antibody/fluorophore pairs was applied to spectrally co-localize them in the same cellular compartment (p-HDAC4^Ser632^/Opal 480, CD31/Opal 480, p16^INK4A^/Opal 520, VCAM1/Opal 570, γ-H2A.X^Ser139^/Opal 690), as recommended by the manufacturer. All multiplex immunofluorescence slides were scanned on a Vectra Polaris Automated Quantitative Pathology Imaging System (Akoya Biosciences) at ×20 magnification, which generates a single unmixed image tiles using Phenochart (Version 1.2.0, Akoya Biosciences). Images were automatically stored on a secure networked server for further analysis.

### Seahorse Mito Stress assay

HAECs were seeded at a density of 5,000 cells/well per on Seahorse XF24 tissue culture plates (Seahorse Bioscience). On the day of experiment, cells were washed and medium was replaced with culture medium supplemented with 25 mM glucose, 2 mM glutamine and 1 mM pyruvate (pH 7.4). For a standard mitochondrial stress test, oligomycin (1 μM), an ATP synthase blocker, FCCP (3 μM), oxidative phosphorylation uncoupler, and Antimycin A/Rotemnone (0.5 μM for each), complex III/ complex I inhibitor, are injected to assess oxygen consumption rate (OCR) and extracellular acidification rate (ECAR) over a 3-min period. At any condition, 5 consecutive measurements of OCR and ECAR are done. Data are normalized to total protein/ well.

### Endothelial cell migration assay

The HAECs were seeded 24 hours before treatment until they become confluent. Cell monolayer migration in response to a cell-free gap created by a sterile 200 μl pipet tip was assessed, as previously described. Images were continuously obtained at 6.4x magnification every 15 min from time point t=0 to t=16 h post-scratch using an incubator-equipped (humidified atmosphere, 37°C, 5% CO_2_) phase-contrast live cell imaging microscope (Olympus IX81) with the Hamamatsu (C11440) detector at 1-megapixel (1024*1024 pixel) 16 bit. The migrated area ratio at 16 h was measured using ImageJ software.

### Endothelial tube formation assay

The angiogenic capacity of endothelial cells represented as the number of tubes formed was examined in HAECs seeded onto Matrigel (Corning^®^ Matrigel^®^-356234) at the sub-confluent level. Images were continuously recorded at 6.4x magnification every 15 min from t=0 to t=16 h using an incubator-equipped phase-contrast Olympus IX81 microscope with the Hamamatsu (C11440) detector at 1-megapixel (1024*1024 pixel) 16 bit, and analyzed using ImageJ Angiogenesis Analyzer software.

### *Ex vivo* aortic ring sprouting assay

Aortic ring sprouting was assessed, as previously described. Fragments of murine aortas (2-mm) were cultured onto Matrigel and subsequently imaged daily (for 5 days) using an incubator-equipped phase-contrast Olympus IX81 microscope, followed by the analysis of the number of aortic sprouts.

### Aortic sprout immunofluorescence staining

Murine aortic endothelial sprouts implanted on Matrigel were fixed in 4% PFA, permeabilized with 0.25% Triton X-100 in PBS and blocked, followed by incubation in primary and secondary antibodies (1:50 to 1:100), respectively. Immunofluorescence imaging was performed by confocal microscopy (Leica TCS-SP8). Nuclei were labeled with the use of DAPI-containing mounting medium for 30 min.

### *Ex vivo* endothelium-dependent relaxation assay

The murine aortas at equal lengths (2 mm) were cut and mounted in a 5-ml organ chamber filled with Krebs-Ringer bicarbonate solution (118.6 mM NaCl, 4.7 mM KCl, 2.5LmM CaCl_2_, 1.2LmM KH_2_PO_4_, 1.2LmM MgSO_4_, 25.1LmM NaHCO_3_, 2.6 μM EDTA, 10.1 mM glucose; 37 °C, pHL7.4), and bubbled with 95% O_2_, 5% CO_2_. Aortic rings were connected to an isometric force transducer (PowerLab 8/30 and LabChart v7.2.5, AD Instruments, Inc.) for continuous isometric tension recording (Multi-Myograph 610 M, Danish Myo Technology, Denmark). Concentration–response curves were obtained in responses to increasing concentrations of acetylcholine (Ach, 10^−9^ to 10^−5^LM; Sigma-Aldrich).

### Statistical analysis

Statistical differences were analyzed by unpaired two-tailed Student’s *t*-test, two-tailed Mann-Whitney *U*-test, and one-way or two-way Analysis of variance (ANOVA) followed by Tukey’s post *hoc* multiple comparison tests. At least three independent triplicated experiments were performed for each experimental set-up. Statistical analysis was performed using GraphPad Prism 8 software (v.8.0.1, La Jolla, CA, USA). Z scores were calculated using R v1.26.1 (Vienna, Austria, 2017) and heatmaps were plotted using the vegan v2.5-5. Spearman rank correlation was used to analyze association between plasma levels of metabolites and differentially-abundant gut microbial taxa. Data are expressed as mean ± SD or SEM, and *P*<0.05 was defined as statistically significant, indicated as *P*<0.05 as *; *P*<0.01 as **; *P*<0.001 as ***; *P*<0.0001 as ****.

### Bioinformatic analysis of gut microbiota data

Bacterial communities were characterized for alpha-diversity using Hill-d0 to 2 (richness) and Pielou’s (evenness) indices. Bacterial community structure (beta-diversity) was also analyzed using Atchinson distance and visualized using Principal Coordinate Analysis (PCoA). Differences were tested using Kruskal-Wallis and PERMANOVA statistical methods for alpha- and beta-diversity, respectively. To analyze significant differences in metagenomic features (including differential abundance of taxa and KO), the Pearson rank correlation analysis was used. The centered log-ratio (CLR) by analysis of composition of microbiomes (ANCOM) was performed to characterize the bacterial taxa harboring VOR, PPFOR, or both gene homologs which are differentially abundant in the microbiome of old *versus* young mice. A Spearman rank correlation was constructed from fecal differentially abundant bacterial taxa harboring VOR or PPFOR homologs and plasma levels of PAA or PAGln measured in the plasma of the corresponding mice using R v1.26.1 (Vienna, Austria, 2017).

Alpha-diversity, beta-diversity, and differential abundance analyses were performed using negative binomial tests as implemented in DESeq2. Multiple comparisons and associated *p*-values were FDR corrected.

## Data availability

The authors declare that all data supporting the findings of this study are available within the paper and its Supplementary information files. The source data underlying Figs. 1b, 1c, 1l, 1m, 2b-e, 3a, 3c-g, 3e, 4b-h, 5a-c, 6a, 6c-i, 6m-p, and Supplementary Figs. 1a-c, 2a, 2b, 3, 4, 5d-i, 6b-d, 7a, and 8b are provided as the Source Data file.

All data from shotgun metagenomics analyses in this paper have been deposited at the Novogene (UK) database (UKPROJ4/XJ/data/Nova_data/1100/220808_A00627_0417_BH32TVDSX5-new), and are publicly available as of request. The data of targeted metabolomics including Thermo’s data files as well as a series of processing information analyzed by Tracefinder software have been deposited in the laboratory’s server. The data from human studies used in this study are held by the Department of Twin Research at King’s College London. The data can be released to bona fide researchers using the normal procedures overseen by the Wellcome Trust and its guidelines as part of our core funding (https://twinsuk.ac.uk/resources-for-researchers/access-our-data/). All other data and reagents that support the findings of this study are available from the lead contacts, Seyed Soheil Saeedi Saravi and Jürg H. Beer, upon request.

## Acknowledgement

We thank Prof. Thomas Michel (Division of Cardiovascular Medicine, Brigham and Women’s Hospital, Harvard Medical School) for helpful discussions and technical support (DAAO constructs). We appreciate Prof. Michael Wannemuehler (Iowa State University) for providing *Clostridium* sp. ASF356.

The authors acknowledge funding from the Swiss National Science Foundation grant #310030_144152, Stiftung Kardio, and Swiss Heart Foundation (to JHB) and from the Novartis Foundation for Medical-Biological Research (#21A053) and the SwissLife Jubiläumsstiftung (#1286) grants (to SSSS). SSSS is also funded by the Fonds zur Förderung des Akademischen Nachwuchses (FAN) and Gebauer Stiftung fellowships.

TwinsUK receives support from grants from the Wellcome Trust (212904/Z/18/Z) and the Medical Research Council (MRC)/British Heart Foundation (BHF) Ancestry and Biological Informative Markers for Stratification of Hypertension (AIM-HY; MR/M016560/1), European Union, Chronic Disease Research Foundation (CDRF), Zoe Global Ltd., the NIHR Clinical Research Facility and Biomedical Research Centre (based at Guy’s and St Thomas’ NHS Foundation Trust in partnership with King’s College London). CM is funded by the Chronic Disease Research Foundation (CDRF) and by the MRC Aim-Hy project grant. IA is funded by Amsterdam Cardiovascular Sciences Post-Doctoral grant (2022-2023).

## Contributions

SSSS and JHB conceptualized and designed the study. SSSS, KG, MA, and PL performed the experiments, including mouse studies, fecal DNA extraction, senescence and signal transduction studies, immunofluorescence assays, angiogenesis assays, and vasorelaxation studies. FC and BP performed shotgun metagenomic data analysis, *in vitro* bacterial studies, and acetate quantification. AT and SL performed LC-MS/MS quantification of plasma PAA and PAGln. CM and IA analyzed and provided data from human TwinsUK cohort. SSSS drafted and wrote the manuscript. SSSS and JHB obtained the grant funding.

## Ethics declarations

### Competing interests

The authors declare no competing interests.

## Supplementary information

## Supplementary Information

**Fig. 1.**
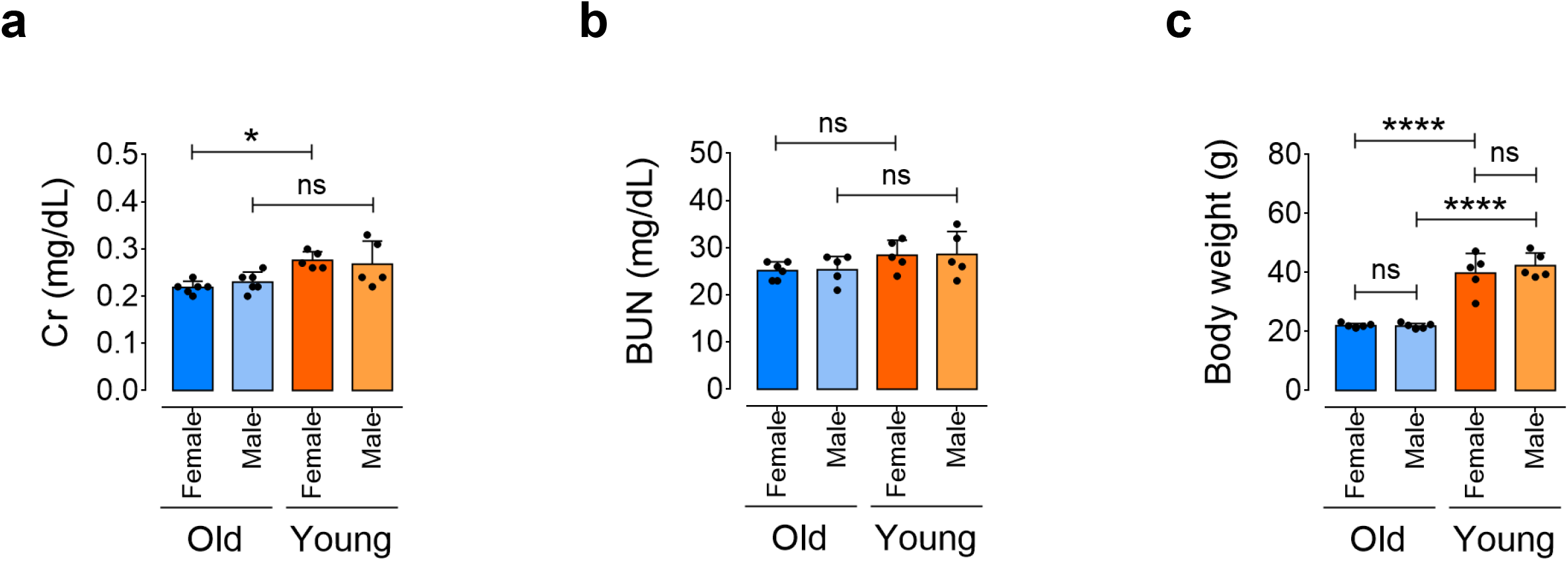
Kidney function profile in aging. **a,** Analysis of body weight in female and male, old and young mice (n=5-6). **b,c,** Bar charts showing kidney function profiles, represented as concentrations of creatinine (Cr) (**b**) and BUN (**c**) in sera of female and male, old and young mice (n=5-6). Error bars represent SD (**a-c**). *P* values were calculated using two-way ANOVA followed by Tukey’s post *hoc* test (**a-c**). (\**P*<0.05, \*\*\*\**P*<0.0001, ns, not significant). Source data are provided as a Source Data file.

**Fig. 2.**
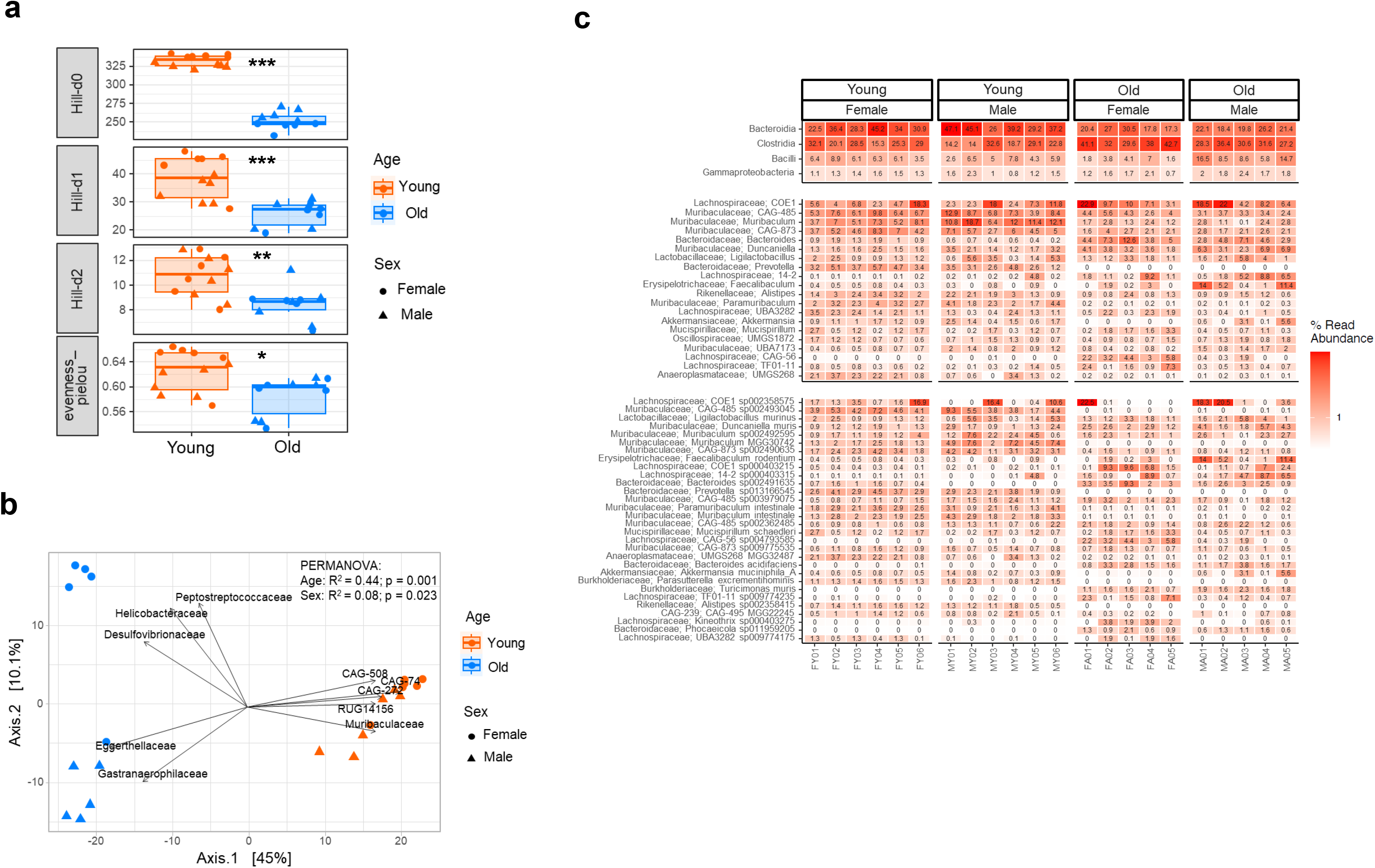
Age-associated gut microbiota alterations. **a,** Box plots representing alpha diversity in feces of both female and male old and young mice measured by Hill-d0 to 2 and Pielou’s evenness indices (n=5-6). **b,** Principal coordinate analysis (PCoA) plot showing beta diversity based on the community level changes in old and young mice (n=5-6). **c,** Heatmap depicting differentially abundant microbial profiles at the strain level (each color represents one bacterial taxa) in the fecal microbiota of female and male, old *versus* young mice (n=5-6). Error bars represent SEM (**a,b**). Data were analyzed using Kruskal-Wallis statistical test (**a**) and Atchinson distance and permutational multivariate analysis of variance (PERMANOVA) tests (**b**). (\**P*<0.05, \*\**P*<0.01, \*\*\**P*<0.001). Source data are provided as a Source Data file.

**Fig. 3.**
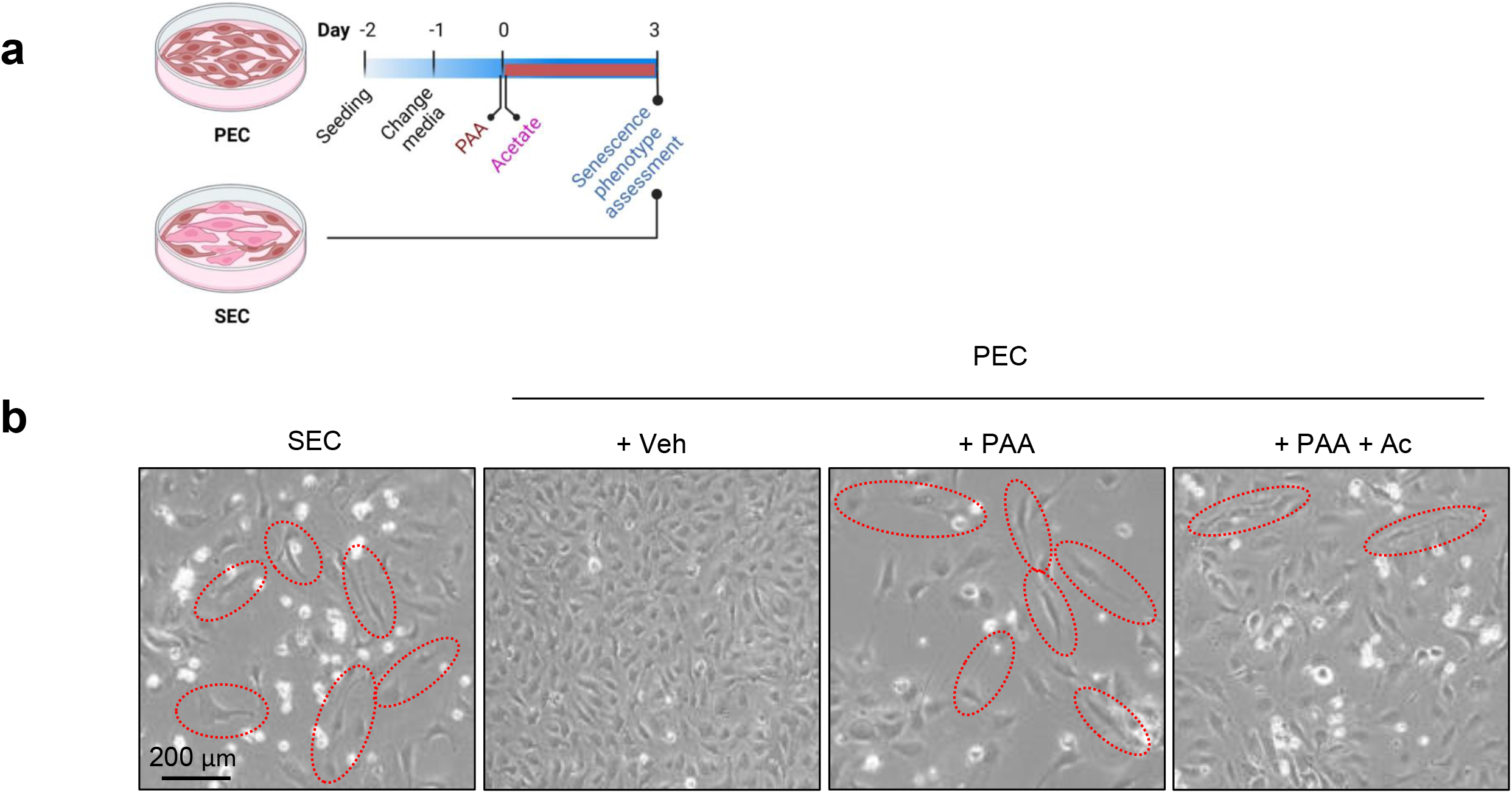
PAA induces senescence phenotype in endothelial cells. **a,** Schematic diagram of the experimental setting: SECs, PECs, PAA-treated PECs (10 μM, for 72 h), and PAA+Acetate-treated PECs (PAA: 10 μM, sodium acetate: 3 μM, for 72 h) were assessed for senescence phenotype. **b,** Representative bright-field images depicting cellular senescence-like phenotype, including enlarged, flattened, and multinucleated appearance, in PECs treated with PAA at the magnitude seen in replicative SECs. The images demonstrate that sodium acetate reduces the number of cells with morphologically senescence-like phenotype in PAA-exposed PECs (n=6). Scale bar, 200 μm. Red circles indicate senescent-like cells. Experiments were triplicated independently.

**Fig. 4.**
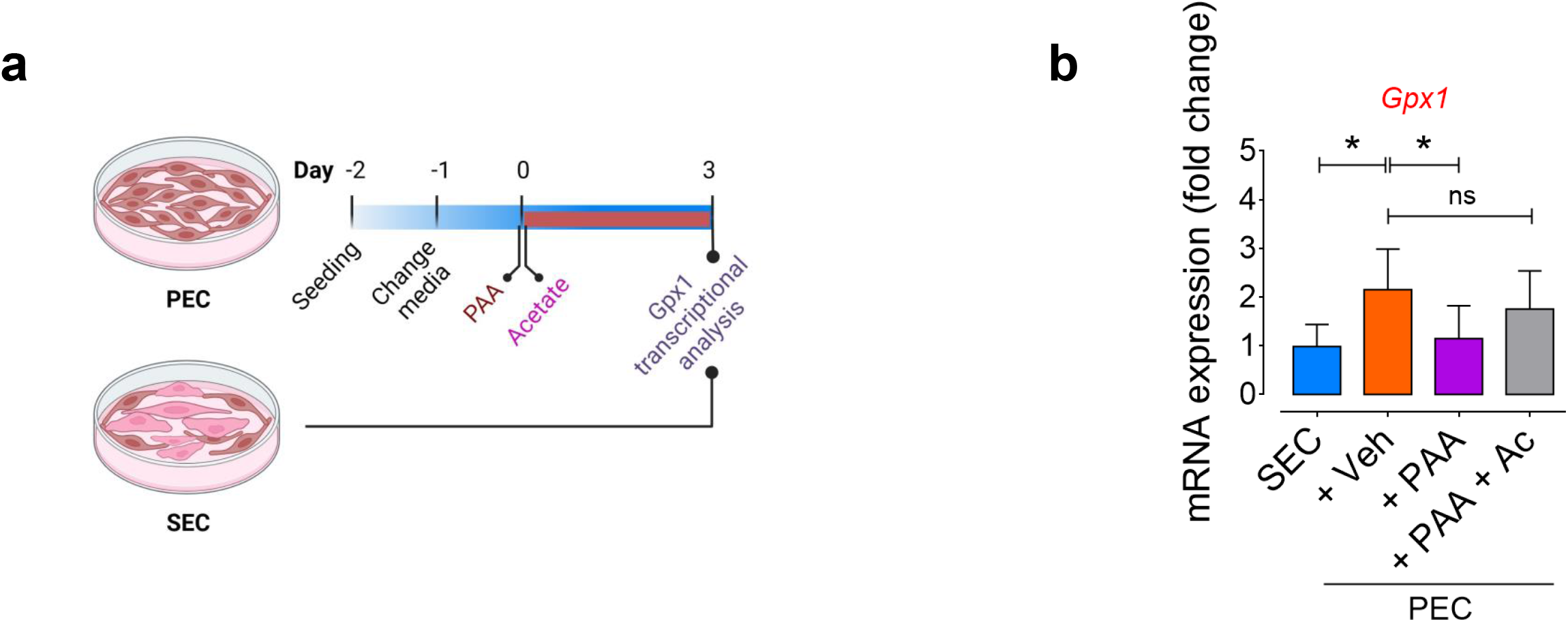
PAA induces oxidative stress by downregulating antioxidant defense in PECs. **a,** Schematic diagram of the experimental setting: SECs, PECs, PAA-treated PECs (10 μM, for 72 h), and PAA+Acetate-treated PECs (PAA: 10 μM, sodium acetate: 3 μM, for 72 h) were subjected to glutathione peroxidase 1 (*Gpx1*) transcriptional analysis. **b,** qPCR analysis demonstrating the downregulation of *Gpx1* in PECs in response to PAA and restoration of the gene expression following the addition of sodium acetate (n=6). Error bars represent SD (**a**). Data represent triplicated biologically independent experiments. *P* values were calculated using one-way ANOVA followed by Tukey’s post *hoc* test (**a-c**). (\**P*<0.05, ns, not significant). Source data are provided as a Source Data file.

**Fig. 5.**
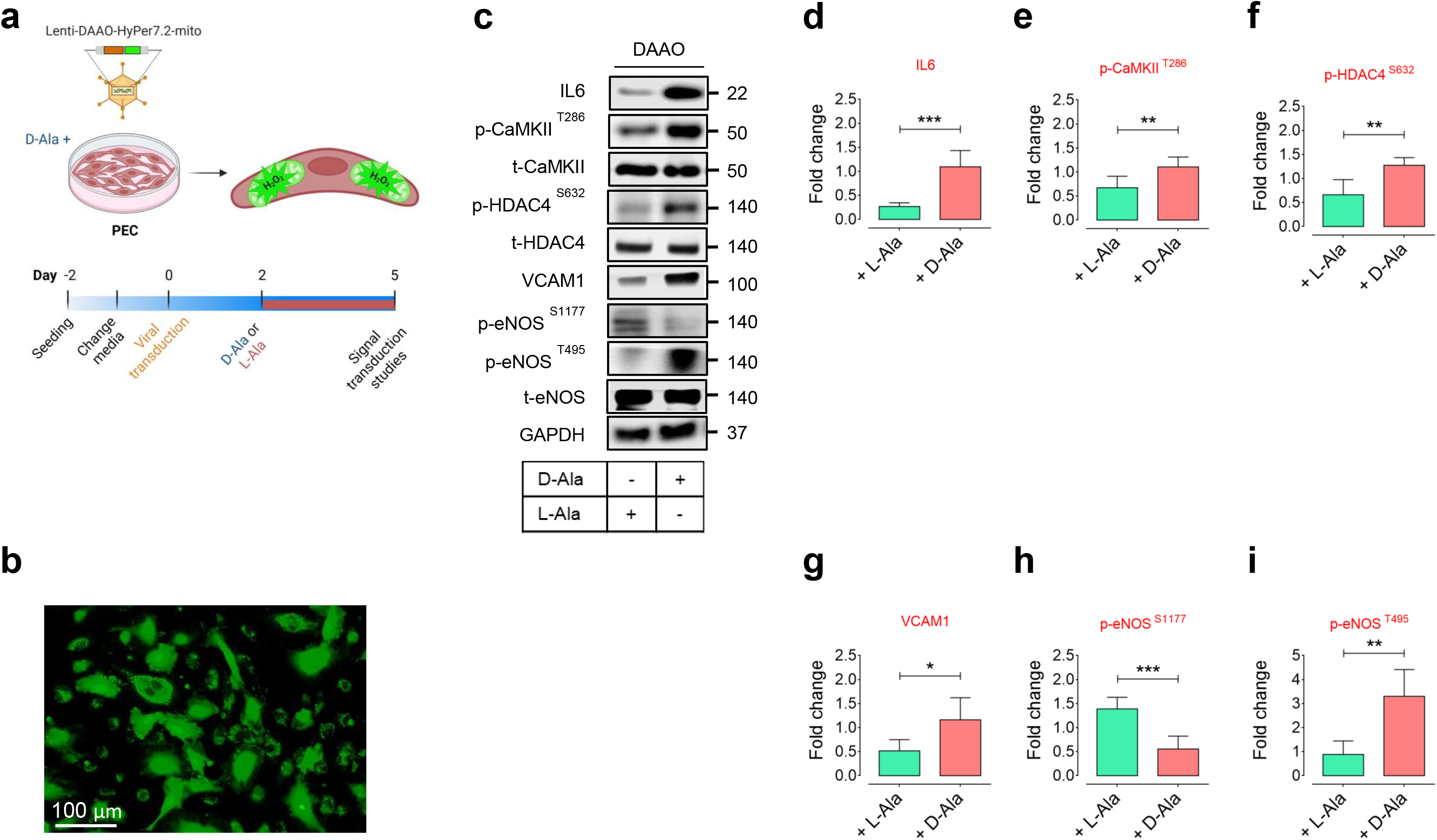
Chemogenetic mitochondrial H_2_O_2_ regulates endothelial NO signaling through IL6-HDAC4 pathway. **a,** Schematic diagram of the experimental setting: PECs were transduced with lentiviral vectors encoding D-aminoacid oxidase (DAAO)-HyPer7.2 targeted to the cell mitochondria for generation of mitochondrial H_2_O_2_ in the presence D-alanine (10 mM) for 72 h. **b,** Representative widefield ratiometric HyPer7.2 images of PECs transduced with Lenti-DAAO-HyPer7.2-mito constructs. **c-i,** Immunoblots (**c**) and quantitative plots show that chemogenetic mitochondrial H_2_O_2_, produced by DAAO in response to D-alanine but not L-alanine, orchestrates IL6-mediated CaMKII-HDAC4 phosphorylation and the subsequent VCAM1-regulated eNOS phosphorylation at Ser1177 and Thr495 in PECs (n=6). Scale bar, 100 μm. Error bars represent SD (**d-i**). Data represent triplicated biologically independent experiments. *P* values were calculated using a two-tailed unpaired Student’s *t*-test (**d-i**). (\**P*<0.05, \*\**P*<0.01, \*\*\**P*<0.001, ns, not significant). Source data are provided as a Source Data file.

**Fig. 6.**
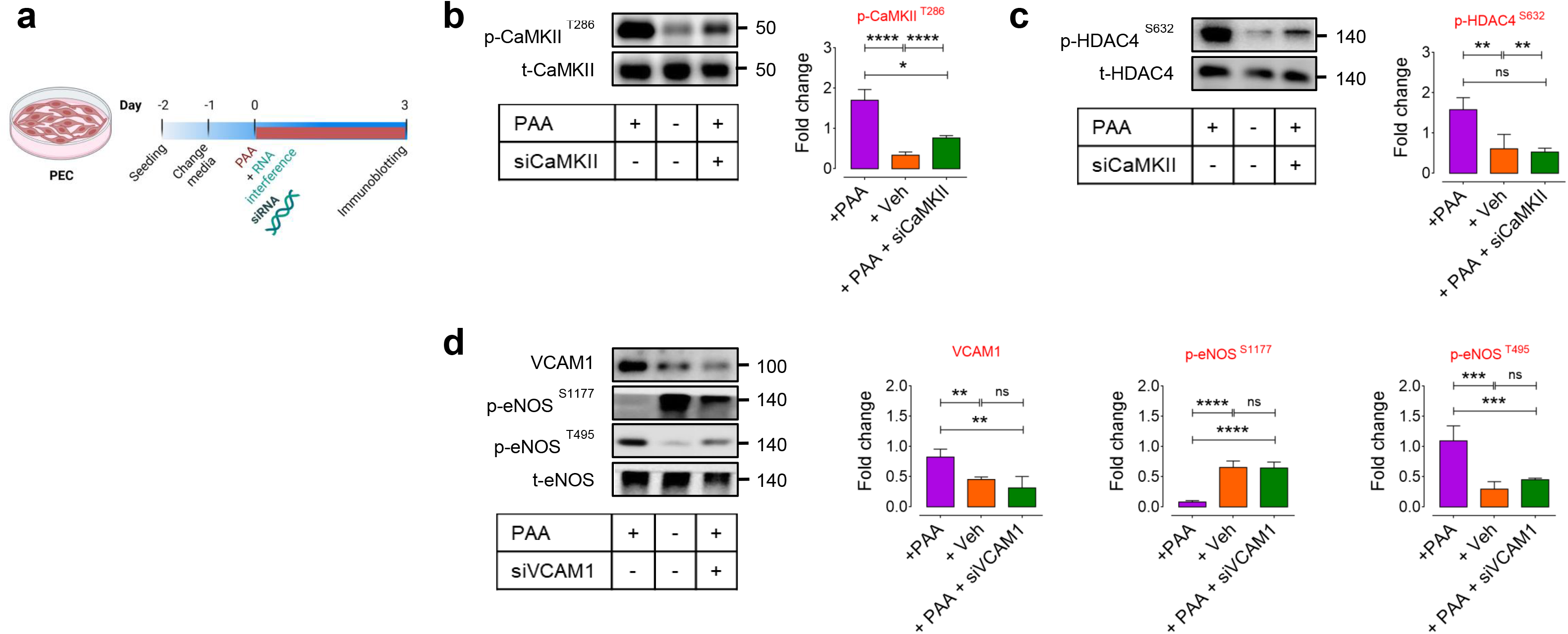
CaMKII-mediated SASP pathway underlies PAA-induced EC dysfunction. **a,** Schematic diagram of the experimental setting: PECs were transfected with siCaMKII or siVCAM1 in the presence or absence of PAA (10 μM). **b,c,** Immunoblots (left) and quantitative plots (right) characterizing the effects of PAA on the PAA-induced post-translational modifications of CaMKII (**b**) and its downstream epigenetic regulator HDAC4 (**c**) using CaMKII knockdown approach in PECs (n=6). **d,** Analysis of VCAM1 expression and eNOS phosphorylation at Ser1177 and Thr495 in siVCAM1-transfected PECs in the presence or absence of PAA (n=6). Error bars represent SD (**b-d**). Data represent triplicated biologically independent experiments. *P* values were calculated using one-way ANOVA followed by Tukey’s post *hoc* test (**b-d**). (\**P*<0.05, \*\**P*<0.01, \*\*\**P*<0.001, \*\*\*\**P*<0.0001). Source data are provided as a Source Data file.

**Fig. 7.**
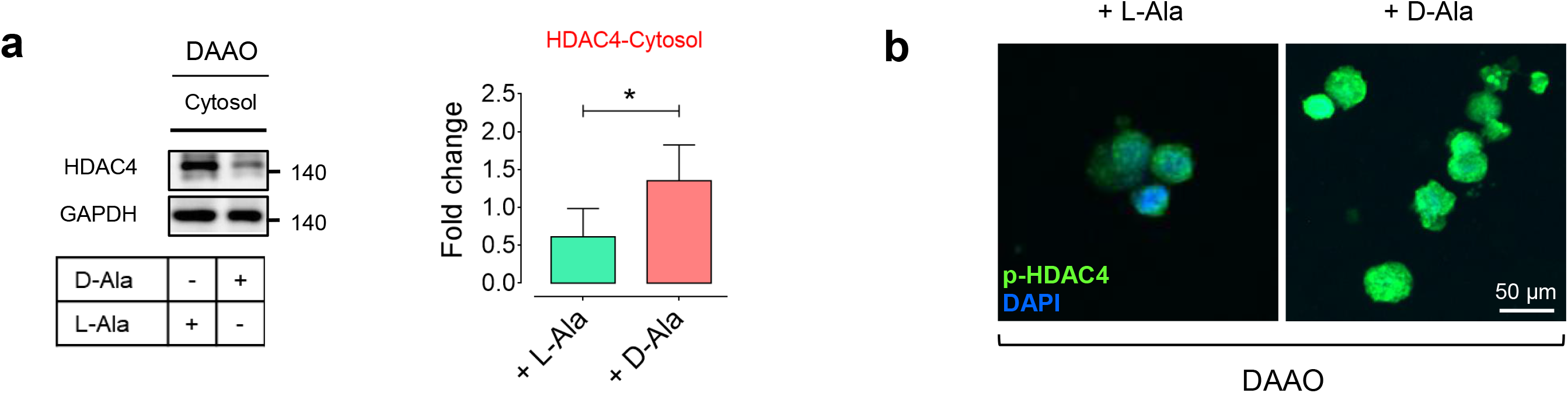
H_2_O_2_ orchestrates HDAC4 phosphorylation and its nuclear export. **a,** Immunoblots (left) and quantitative plot (right) reveal HDAC4 unclear export, represented as an increased expression in the cytosol in response to intracellular H_2_O_2_ generated by DAAO in the presence of D-alanine (10 mM, for 72 h) in PECs (n=6). **b,** p-HDAC4 immunostaining shows that intracellular H_2_O_2_ markedly increases HDAC4 phosphorylation that facilitates its translocation towards the cell cytosol in PECs (n=6). Scale bar, 50 μm. Error bars represent SD (**a**). Data represent triplicated biologically independent experiments. *P* values were calculated using a two-tailed unpaired Student’s *t*-test (**a**). (\**P*<0.05, \*\**P*<0.01, \*\*\**P*<0.001, ns, not significant). Source data are provided as a Source Data file.

**Fig. 8.**
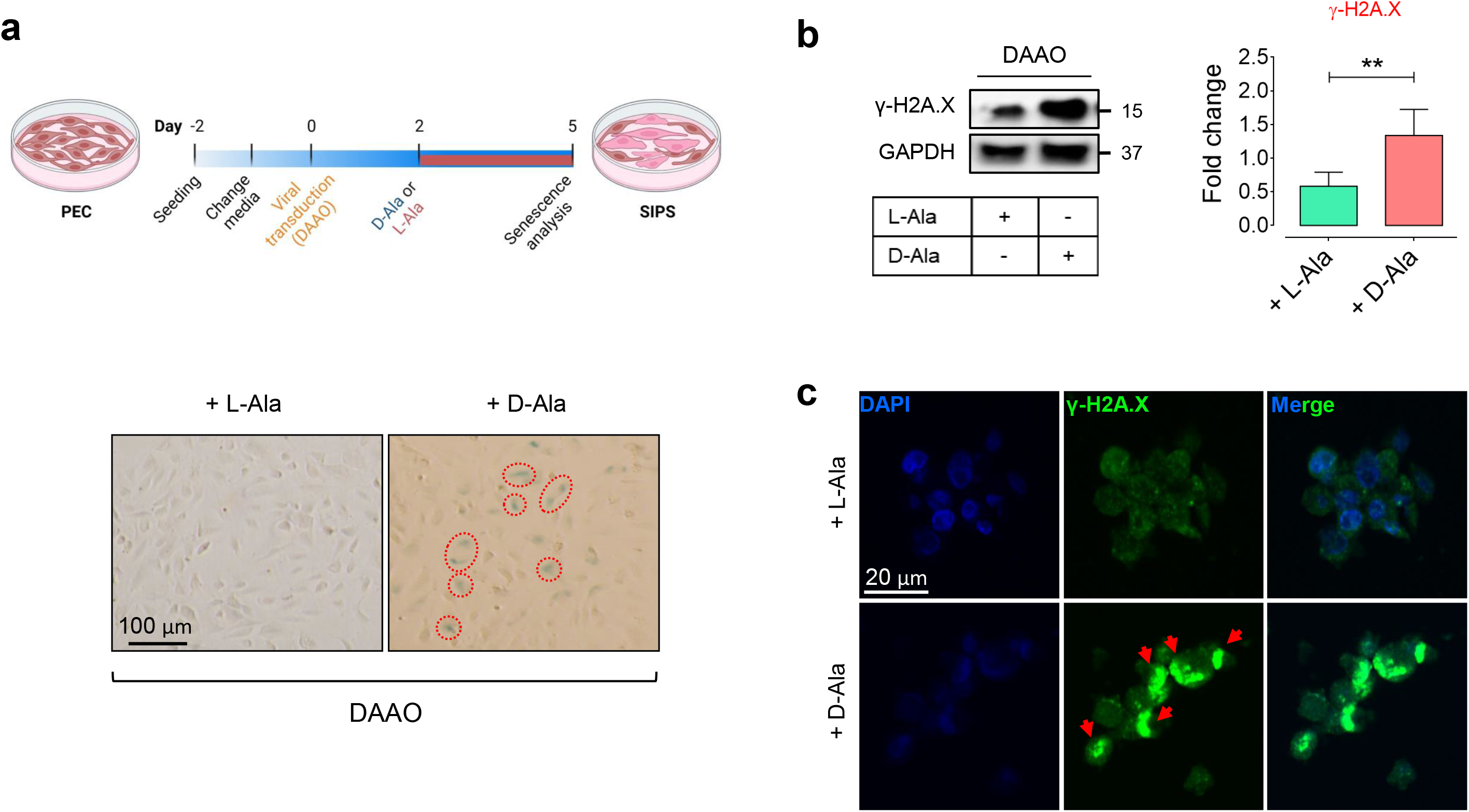
Chemogenetic H_2_O_2_ promotes EC senescence. **a,** Schematic diagram of the experimental setting: PECs were transduced with lentiviral vectors encoding D-aminoacid oxidase (DAAO) targeted to the cell mitochondria for generation of mitochondrial H_2_O_2_ in the presence D-alanine (10 mM) for 72 h. Representative images of SA-β-gal staining demonstrates that DAAO-mediated mitochondrial H_2_O_2_ induced cellular senescence in PECs, as shown by increased SA-β-gal-positive cells (%) (n=6). **b,c,** Immunoblots (**b**) and γ-H2A.X immunostaining images (**c**) reveal a marked increase in DDR following mitochondrial H_2_O_2_ production in DAAO-transduced PECs incubated with D-alanine (n=6). Scale bar, 20 and 100 μm. Red circles indicate SA-β-gal-positive cells. Data represent triplicated biologically independent experiments. Source data are provided as a Source Data file.

**Fig. 9.**
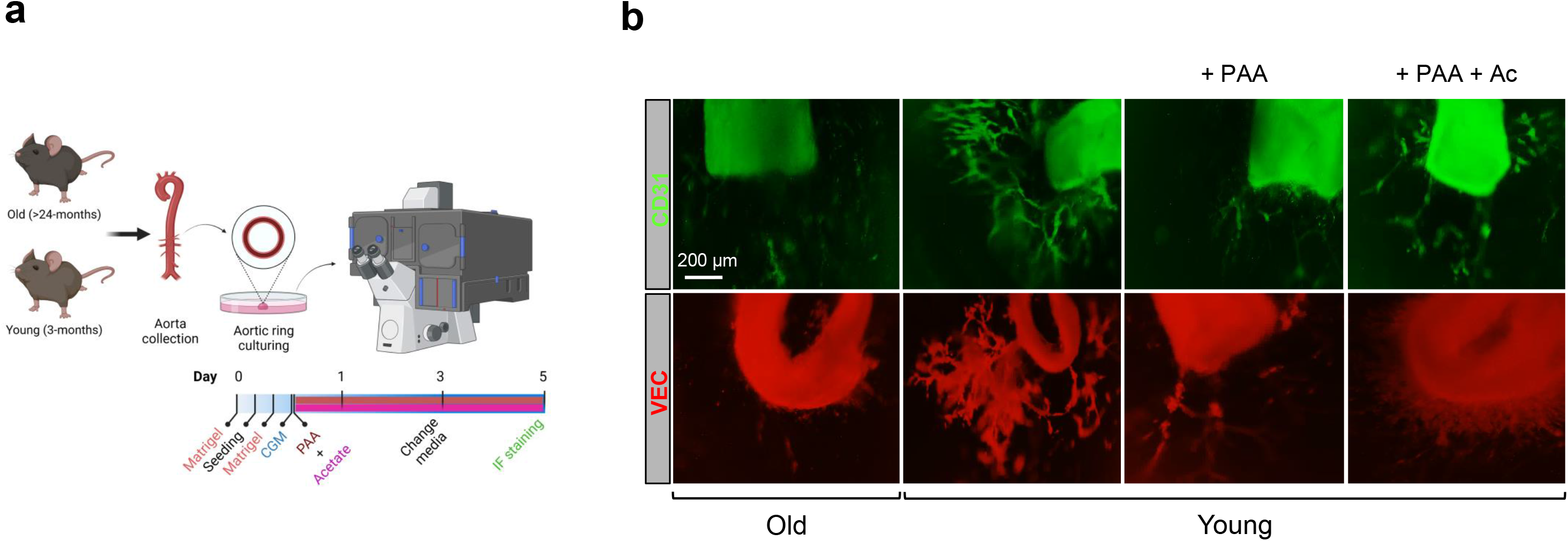
PAA reduces aortic endothelial sprouting. **a,** Schematic diagram of the experimental setting: aortic rings from old and young mice were subjected to PAA (10 μM) for 72 h and tested for endothelial sprouting, followed by CD31 (green) and VEC (red) immunofluorescence staining. **b,** CD31 and VEC immunofluorescence staining in mouse aortas showing decreased endothelial sprouting after incubation of young aortic rings with PAA as opposed to vehicle. Immunofluorescence images reveal that co-treatment with sodium acetate markedly restores angiogenic capacity of aortic CD31, VEC-positive ECs (n=6). Scale bar, 200 μm. Experiments were triplicated independently.

**Supplementary Table 1.**
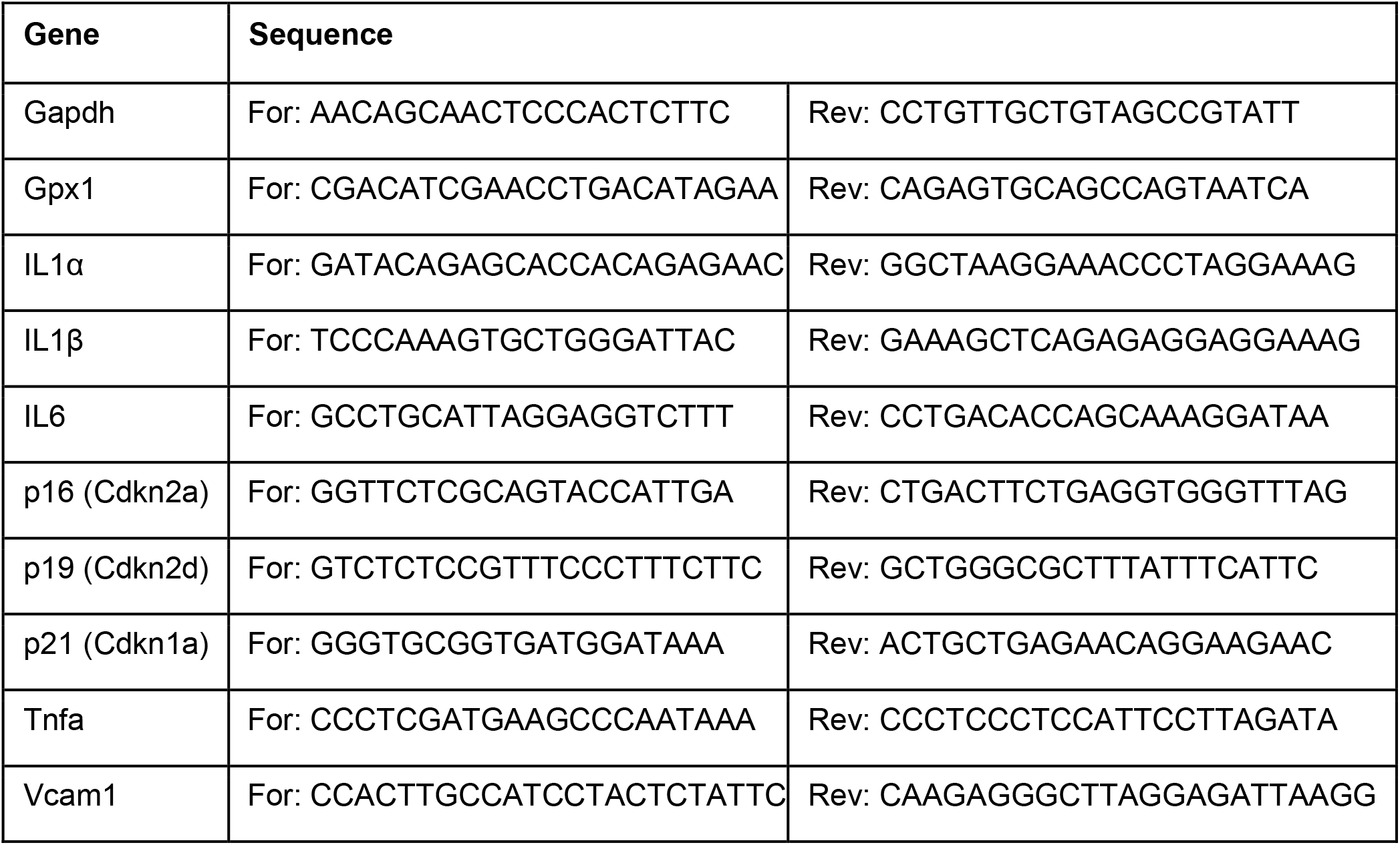
Gene-specific forward and reverse primers for RT-PCR amplification. *Gapdh*, Glyceraldehyde 3-phosphate dehydrogenase; *Gpx1*, Glutathione peroxidase 1; IL1α, Interleukin 1-alpha; *IL1β*, Interleukin 1-beta; *IL6*, Interleukin 6; *p16*, Cdkn2a; *p19*, Cdkn2d; *p21*, Cdkn1a; *Tnfa*, tumor necrosis factor-α; *Vcam1*, Vascular cell adhesion molecule 1.

## Notes

### Competing Interest Statement

The authors have declared no competing interest.

